# Reference genomes of four miniature and non-miniature cypriniform fishes inhabiting acidic peat-swamp forest blackwaters of Southeast Asia

**DOI:** 10.64898/2026.03.21.713365

**Authors:** Hiranya Sudasinghe, Zuyao Liu, Laura Triginer-Llabrés, Heok Hui Tan, Ralf Britz, Walter Salzburger, Catherine L. Peichel, Lukas Rüber

## Abstract

The acidic blackwaters of Southeast Asia’s peat-swamp forests represent some of the most extreme freshwater environments on Earth. Despite their very low pH values, limited nutrients, and hypoxic conditions, these blackwater habitats harbor a remarkable diversity of freshwater fishes, including multiple lineages that have independently adapted to these extreme conditions and, in some cases, exhibiting extreme body miniaturization. These replicate evolutionary lineages therefore provide a powerful comparative framework to investigate adaptation to extreme environments and the genomic basis of miniaturization. Here, we present high-quality, annotated reference genomes for four cypriniform species endemic to these peat-swamp forest ecosystems: *Paedocypris* sp., *Sundadanio atomus*, *Boraras brigittae*, and *Rasbora kalochroma*. The first two are progenetic miniatures, including *Paedocypris*, comprising the smallest known fish, while *B. brigittae* represents a proportioned dwarf and *R. kalochroma* a non-miniature taxon. Genome sizes ranged from 401-1,290 Mb and heterozygosity from 0.34-1.7%. All genome assemblies achieved pseudo-chromosome-level contiguity, high k-mer completeness (>99%), and high BUSCO completeness (94.5-98.9%). Repeat analyses revealed lineage-specific differences in transposable element landscapes and abundances, while gene annotation identified notable intron length reduction in progenetic miniatures.

## Introduction

Extreme freshwater environments are typically characterized by low productivity, reduced biodiversity, and strong abiotic constraints that impose multiple challenges to the residing organisms with respect to physiology, energy metabolism, and life-history strategies (Bell & Callaghan, 2012; Plath et al., 2015; Rothschild & Mancinelli, 2001). Although such environments often harbor highly specialized taxa, they commonly lack many ecosystem processes typical of more benign habitats, including primary production and nitrogen fixation. Only a small number of freshwater fishes occupy such extreme habitats, yet those that do frequently exhibit striking physiological and morphological adaptations that enable persistence under challenging conditions.

Freshwaters with a pH < 4 are considered extremely acidic and pose multiple stressors due to increased metal solubility, ionoregulatory disruptions, and oxidative stress (Rothschild & Mancinelli, 2001; Spijkerman & Weitholf, 2012). For fishes, chronic exposure to acidic conditions affects ion balance, respiration, reproduction, and early development, making these environments particularly demanding (Nelson, 2015). Among the world’s freshwater ecosystems, the tannin-rich blackwaters of Southeast Asia’s peat swamp forests (Fig. 1a) are among the most chemically extreme. These peatlands, globally significant both for biodiversity and carbon sequestration, are highly acidic (pH 3.3–5.9), hypoxic, and enriched in dissolved organic carbon (Dommain et al., 2014; Omar et al., 2022; Page et al., 2022; Posa et al., 2011). Despite these constraints, they support an unexpectedly rich and largely endemic fish fauna, much of which has been described only in recent decades (Beamish et al., 2003; Kottelat et al., 2006; Ng et al., 1994; Thornton et al., 2018). Many of these species occur exclusively in acidic blackwaters, underscoring their ecological specialization.

**Fig. 1.**
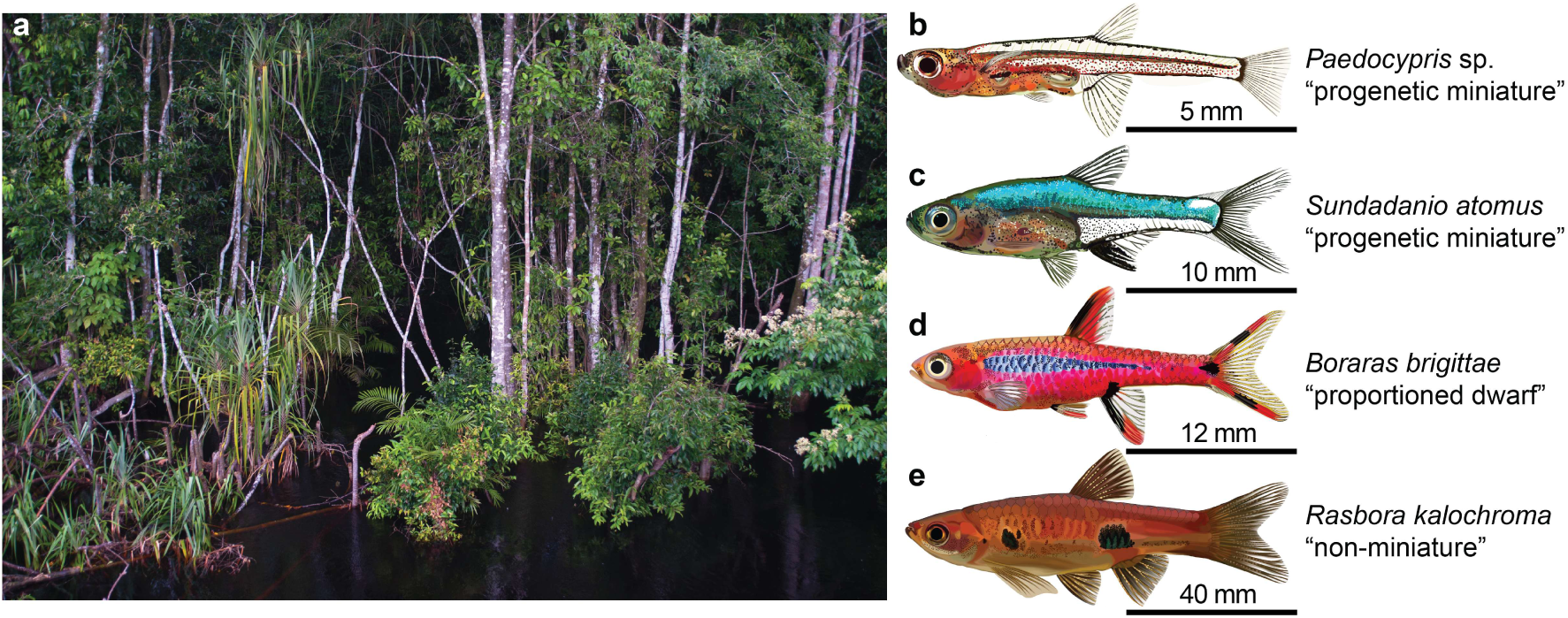
**(a)** Blackwater peat swamp forest habitat in Southeast Asia, an extreme environment supporting a unique assemblage of freshwater fishes. Panels **b–e** shows the species for which high-quality reference genome assemblies were generated in this study: **(b)** *Paedocypris* sp.; **(c)** *Sundadanio atomus*; **(d)** *Boraras brigittae*; and **(e)** *Rasbora kalochroma*.

A distinctive feature of these fish assemblages is the prevalence of miniature species, defined as those maturing at ≤ 26 mm standard length and often exhibiting paedomorphic skeletal and morphological traits (Kottelat & Vidthayanon, 1993; Weitzman & Vari, 1988). Miniaturization has evolved repeatedly within Cypriniformes, the most species-rich order of freshwater fishes. This order contains both “proportioned dwarfs,” which resemble scaled-down versions of their larger relatives, and “progenetic miniatures,” which display extreme developmental truncation and retain larval features into adulthood (Britz et al., 2014; Britz & Conway, 2009; Rüber et al., 2007). Interestingly, progenetic miniatures such as *Paedocypris*, including *P. progenetica*, the smallest known freshwater fish maturing at 7.9 mm SL (Kottelat et al., 2006), co-occur alongside proportioned dwarfs and non-miniature relatives in peat swamp forests (Fig. 1b). Miniaturization in these systems may reflect adaptation to chronically low-resource, acidic conditions.

The coexistence of multiple independent lineages that have repeatedly adapted to the same extreme environmental pressures makes the fish assemblages of peat-swamp forest an ideal natural experiment for studying convergent evolution, ecological specialization, and the genomic mechanisms underlying extreme miniaturization (Malmstrøm et al., 2018). Yet, despite their ecological and evolutionary significance, the genomic basis of adaptation to acidic blackwaters remains largely unexplored. Moreover, peat-swamp forest ecosystems are among the most threatened habitats in Southeast Asia, and the lack of genomic resources could limit efforts to assess evolutionary distinctiveness, population resilience, and adaptive potential in these highly specialized species (Giam et al., 2012; Theissinger et al., 2023).

Here, we report the generation of four high-quality reference genomes representing independent lineages of cypriniform fishes inhabiting the extreme peat-swamp forest environments of Southeast Asia. These include two progenetic miniatures (*Paedocypris* sp. “Kalimantan Tengah” and *Sundadanio atomus*), one proportioned dwarf (*Boraras brigittae*), and one non-miniature species (*Rasbora kalochroma*). Together, these assemblies provide valuable genomic resources to enable future comparative analyses of adaptation to extreme environments, the evolution of miniaturization, and the genomic consequences of developmental truncation. They also offer important data for conservation genomics in one of the world’s most threatened and least studied freshwater ecosystems.

## Materials and methods

### Taxon sampling

We selected four species of Cypriniformes endemic to peat swamp forest blackwaters in Southeast Asia (Supplementary Table 1; Fig. 1). These included two progenetic miniature species (species that exhibit truncated development and retain larval characteristics resembling an early developmental stage of their larger relatives): *Paedocypris* sp. “Kalimantan Tengah” (hereafter *Paedocypris* sp.) and *Sundadanio atomus*; one proportioned dwarf (small-bodied species morphologically similar to their larger relatives): *Boraras brigittae*; and one non-miniature species: *Rasbora kalochroma*. Specimens were acquired through the aquarium trade in Singapore. Individuals were euthanised using an overdose of MS-222, flash-frozen in liquid nitrogen, and stored at −80 °C at the Lee Kong Chian Natural History Museum, National University of Singapore, prior to export to the Naturhistorisches Museum Bern, Switzerland.

### DNA extraction, library preparation, and sequencing

For PacBio HiFi sequencing, high molecular weight (HMW) genomic DNA was extracted from multiple tissues of a single individual per species using the Qiagen Genomic-tips 20/G columns (Qiagen, 10223) with the Genomic DNA Buffer Set (Qiagen, 19060), following the instructions for tissue genomic DNA (gDNA) extraction. DNA quality and integrity were assessed using a Qubit 4.0 fluorometer (Thermo Fisher Scientific), an advanced analytical FEMTO Pulse instrument, and a Denovix DS-11 UV-Vis spectrophotometer.

Between 1000-5000 ng of gDNA was used to prepare barcoded PacBio SMRTbell libraries for each species. A Megaruptor 3 was used to shear the gDNA, which was then concentrated and cleaned using 1 x SMRTbell clean-up beads. For *Paedocypris* sp., an insert size of 6–7 kb was obtained, and sequencing was carried out on a PacBio Sequel IIe platform using a Single-Molecule Real-Time (SMRT) cell. For the other three species, insert sizes of 10–12 kb were achieved, and these libraries were pooled and sequenced on a PacBio Revio system containing 4 x SMRT cell.

To support chromosome-level scaffolding, Hi-C libraries were prepared from muscle tissue of a different individual for each species using the Arima High Coverage Hi-C Kit, following the manufacturer’s protocol for low input, flash frozen animal tissue. For this protocol, 2-50 mg of fish muscle tissue were used as input and 12 PCR cycles were employed in the Hi-C library preparation. Sequencing was performed on an Illumina NovaSeq 6000 system with paired-end 150 bp reads.

For transcriptome-based genome annotation, full-length isoform sequencing (Iso-Seq) libraries were generated from RNA extracted from whole-body tissues using the Direct-zol RNA Miniprep kit (Zymo Research, R2051) according to the kit instruction manual for animal tissues and with the on-column DNaseI treatment. The quantity and quality of the purified total RNA was assessed using a Qubit 4.0 fluorometer with the Qubit RNA BR Assay Kit (Thermo Fisher Scientific, Q10211) and an Advanced Analytical Fragment Analyzer System using a Fragment Analyzer RNA Kit (Agilent, DNF-471). For *Paedocypris* sp., RNA was pooled from two individuals; for the other three species, RNA was extracted from a single individual per species. PacBio IsoSeq v2 libraries were prepared starting with 300 ng of RNA as input and sequenced on a PacBio Sequel IIe platform. Thereafter, the CCS generation was performed on the Sequel IIe instrument and the barcode demultiplexing workflow as well as IsoSeq pipeline was run in SMRT Link v12.

All DNA and RNA extractions, library preparations, and sequencing were conducted at the Next Generation Sequencing (NGS) Platform, University of Bern, Switzerland.

### Reference genome assembly

Our genome assembly pipeline largely followed the Vertebrate Genome Project (VGP) standard workflow as described in Larivière et al. (2024), with several modifications at specific stages.

As a first step, HiFiAdapterFilt (Sim et al., 2022) was used to remove potential adapter sequences and other artifacts from the PacBio HiFi reads. To estimate genome size, heterozygosity, and repetitive content, we performed k-mer analysis using Meryl (Rhie et al., 2020) and GenomeScope2.0 (Ranallo-Benavidez et al., 2020) with k=31.

Since the HiFi and Hi-C reads originated from two different individuals for each species, we used Hifiasm (Cheng et al., 2021) in solo mode with light purge settings, using only HiFi reads. This generated a pseudohaplotype assembly consisting of a primary and an alternate contig set. The primary contigs were then processed with purge_dups (Guan et al., 2020) to identify and remove haplotypic duplication. Next, the alternate contigs were concatenated with these haplotigs from the purged primary contigs, and purge_dups was run again to further refine the alternate assembly. This produced a final purged primary and alternate assembly.

Mitochondrial genome assembly was performed using MitoHiFi (Uliano-Silva et al., 2023) on the purged primary assembly. Reference mitogenomes were used for species-specific assembly: NC_020436.1 (*Paedocypris progenetica*), AP011431.1 (*Sundadanio axelrodi* ’blue’), AP011420.1 (*Boraras maculatus*), and NC_063882.1 (*Rasbora kalochroma*) for *Paedocypris* sp., *Sundadanio atomus*, *Boraras brigittae*, and *Rasbora kalochroma*, respectively. Assembled mitogenomes were aligned to the genome assembly using BLASTn (Camacho et al., 2009) to locate scaffolds containing mitochondrial sequences.

Hi-C data preprocessing was performed using fastp (Chen et al., 2018) in two stages. In the first stage, adapter sequences, polyG/polyX tails, and duplicated reads were removed. In the second stage, quality filtering was applied using a sliding window approach with a minimum Phred score of 20. The cleaned Hi-C reads were then aligned to the purged primary assembly using Chromap (Zhang et al., 2021). SAMtools (Li et al., 2009) was used to convert SAM to BAM format and to index the genome.

Scaffolding was carried out using YaHS with default parameters (Zhou et al., 2023) with the purged primary assembly and Hi-C reads. To identify potential contamination, the scaffolded assembly was soft-masked with Dustmasker (Morgulis et al., 2006) and screened using Kraken2 (Wood et al., 2019) against a database comprising archaeal, bacterial, plasmid, viral, and UniVec_Core sequences. Contaminant scaffolds identified by Kraken2 were flagged for removal. Furthermore, the assemblies were screened for contamination using the BlobToolKit environment (Challis et al., 2020). The genome coverage needed for the BlobToolKit was calculated using Mosdepth (Pedersen & Quinlan, 2018).

All contaminant scaffolds (from Kraken2 and BlobToolKit) and mitochondrial scaffolds (identified via BLAST) were removed from the assembly using SeqKit2 (Shen et al., 2024), yielding a decontaminated nuclear genome.

These decontaminated genome assemblies were then re-scaffolded with Chromap and YaHS. We used the auxiliary juicer-pre tool (available in the YaHS pipeline) and Juicer Tools to generate a Hi-C contact maps. These maps were manually reviewed and curated using Juicebox (Durand et al., 2016). The curated contact maps were finalized using juicer-post. We then used TGS-GapCloser (Xu et al., 2020) to close sequence gaps in the assemblies. Using seqkit, scaffolds in this pseudo-chromosomal assemblies were sorted by length and renamed, assigning the longest scaffold to Scaffold_1 and numbering the remaining scaffolds in decreasing size.

Assembly quality were assessed at multiple stages using: Merqury (Rhie et al., 2020) for consensus QV and k-mer completeness; and gfastats (Formenti et al., 2022) for genome assembly statistics; and benchmarking sets of universal single-copy orthologs, BUSCO (Manni et al., 2021), reimplemented in Compleasm (Huang & Li, 2023), using the actinopterygii_odb10 (n=3640) lineage dataset for genome completeness.

To assess overall genome architecture and synteny, we generated dot plots using the web-based D-GENIES tool (Cabanettes & Klopp, 2018) to compare our assemblies with the *Danio rerio* GRCz11 reference genome, enabling visualization of large-scale structural patterns and conserved chromosomal blocks.

### Genome annotation

Extensive de novo TE Annotator (EDTA) was employed to construct a species-specific transposable element (TE) library for each genome assembly (Ou et al., 2019). EDTA was run with the --sensitive 1 option to enable RepeatModeler-based identification of additional TEs (Flynn et al., 2020), and with --anno 1 to perform whole-genome TE annotation following library construction. The resulting custom repeat libraries for each genome were subsequently used to perform genome-wide repeat masking in RepeatMasker with a slow search which is more sensitive (Smit et al., 2015). For each species, the CpG-adjusted Kimura divergence from the consensus repeat sequences was estimated using the RepeatMasker script calcDivergenceFromAlign.pl, applied to the .align files produced during genome masking. The generated .divsum files were imported into R Statistical Software 4.3.3 (R Core Team, 2024), processed with custom code, and plotted using the ggplot2 package (Wickham, 2016).

Gene annotation was performed using the BRAKER3 pipeline with Iso-Seq long-read transcriptomic data (Brůna et al., 2024; Gabriel et al., 2024). Following the workflow of Card et al. (2019), complex repeat annotations were separated from low-complexity repeats, and the resulting complex-repeat GFF file was used to generate both soft-masked and hard-masked versions of each genome assembly using bedtools (Quinlan & Hall, 2010). Iso-Seq reads were aligned to the hard-masked genome using minimap2 (Li, 2018), and the resulting SAM files were converted to sorted and indexed BAM files with samtools. For protein homology evidence, we used the Vertebrata protein set from OrthoDB v11 (Kuznetsov et al., 2023). The soft-masked assembly, the Iso-Seq BAM file, and the protein database were provided as inputs to BRAKER3. Within the pipeline, StringTie2 (Kovaka et al., 2019) was used to assemble a draft transcriptome, which served as the basis for transcript-supported gene prediction with GeneMarkS-T (Tang et al., 2015). BRAKER3 then used GeneMark-ETP (Brůna et al., 2024) to integrate transcript, protein, and ab initio evidence and to select gene models with high similarity scores and high confidence. GeneMark-ETP generated three categories of extrinsic hints using ProtHint (Brůna et al., 2020): (a) hints supported by both transcript and protein alignments, (b) hints supported by transcript evidence and ab initio predictions, and (c) hints supported solely by protein similarity. These hints were combined to construct a set of high-confidence seed genes. AUGUSTUS (Stanke et al., 2006, 2008) was trained on this high-confidence gene set and used to perform ab initio genome-wide gene prediction. The outputs from AUGUSTUS and GeneMark-ETP were then integrated using TSEBRA (Gabriel et al., 2021) to obtain the final, non-redundant set of high-confidence protein-coding genes. BRAKER3 was executed in BUSCO-completeness-maximizing mode, running compleasm with the actinopterygii_odb10 lineage in genome mode and converting the identified BUSCO matches into additional hints for AUGUSTUS. All analyses were conducted within the BRAKER3 Singularity container (braker3_lr.sif).

The completeness of the BRAKER3 annotations was assessed using BUSCO in protein mode with the actinopterygii_odb10 lineage. We additionally evaluated proteome completeness and orthology consistency using OMArk (Nevers et al., 2025). OMArk assesses proteome quality by comparing the target proteome to an inferred ancestral gene repertoire for the chosen lineage, reconstructed from extant species. This analysis reports both completeness and consistency based on lineage-appropriate gene family assignments. To run OMArk, we first generated an OMAmer database for each predicted proteome using the LUCA.h5 reference (Rossier et al., 2021), and the resulting OMAmer search output was then provided as input to OMArk. Both BUSCO and OMArk were run on the longest isoforms which were filtered from the script agat_sp_keep_longest_isoform.pl available in AGAT (Dainat, 2022).

For *Rasbora kalochroma* and *Paedocypris sp*., OMArk initially reported a substantial number of “unknown” or “misplaced” genes, prompting additional curation steps to remove false-positive gene predictions. For these “unknown” or “misplaced” genes, we quantified overlap between predicted coding sequences (CDS) and repeat annotations and flagged CDS with >50% overlap. We then performed tblastn searches of these predicted proteins against the EDTA-derived TE library to identify gene models with high similarity to transposable elements. In addition, we extracted proteins that met any of the following criteria: zero mean Iso-Seq coverage, length <150 amino acids, or single-exon structure. Gene models meeting these criteria are commonly associated with transposable-element fragments, pseudogenes, or assembly artefacts, and are therefore likely to represent false-positive predictions and were removed from the final annotations (Caballero & Wegrzyn, 2019; Ou & Jiang, 2018; Vuruputoor et al., 2023).

Summary statistics for the final BRAKER3 annotations were calculated using agat_sp_statistics.pl in AGAT. The list of software versions used, and their sources is given in Supplementary Table 2.

## Results and discussion

### Genome size and heterozygosity estimation

K-mer based genome size estimates for the four species ranged from 401.5 to 1290.8 Mb (Fig. 2; Supplementary Table 3). The two progenetic miniatures, *Paedocypris* sp. and *Sundadanio atomus*, exhibited the smallest genomes at 401.5 Mb and 654.5 Mb, respectively, whereas the proportioned dwarf *Boraras brigittae* had an intermediate genome size of 989.8 Mb, and the non-miniature *Rasbora kalochroma* possessed the largest genome at 1290.8 Mb. The genome size estimated for *Paedocypris* sp. is comparable to that reported for *P. micromegethes* (412.6 Mb) and slightly lower than the estimate for *P. carbunculus* (437.3 Mb) (Malmstrøm et al., 2018). The genome size of *R. kalochroma* is consistent with that of *R. borapatensis* (C-value = 1.44 pg; ∼1408.3 Mb) but is substantially larger than that of *R. dandia* (C-value = 0.85 pg; ∼831.3 Mb) reported in the Animal Genome Size Database (Gregory, 2025). No comparable genome size estimates are currently available for either *Boraras* or *Sundadanio*. The bimodal coverage distribution in the k-mer plots in all four species is indicative of diploid genomes, with the first (heterozygous) peak in the range of 21-30-fold coverage and the second (homozygous) peak in the range of 42-60-fold coverage (Fig. 2). Estimated heterozygosity values ranged from 0.34% to 1.7%, with the lowest observed in *B. brigittae* and the highest in *Paedocypris* sp. (Fig. 2; Supplementary Table 3). Notably, the 1.7% heterozygosity in *Paedocypris* sp. characterized with a strong first peak in the k-mer plot (Fig. 2a) is relatively high compared with other diploid cypriniform fishes (Oriowo et al., 2025).

**Fig. 2.**
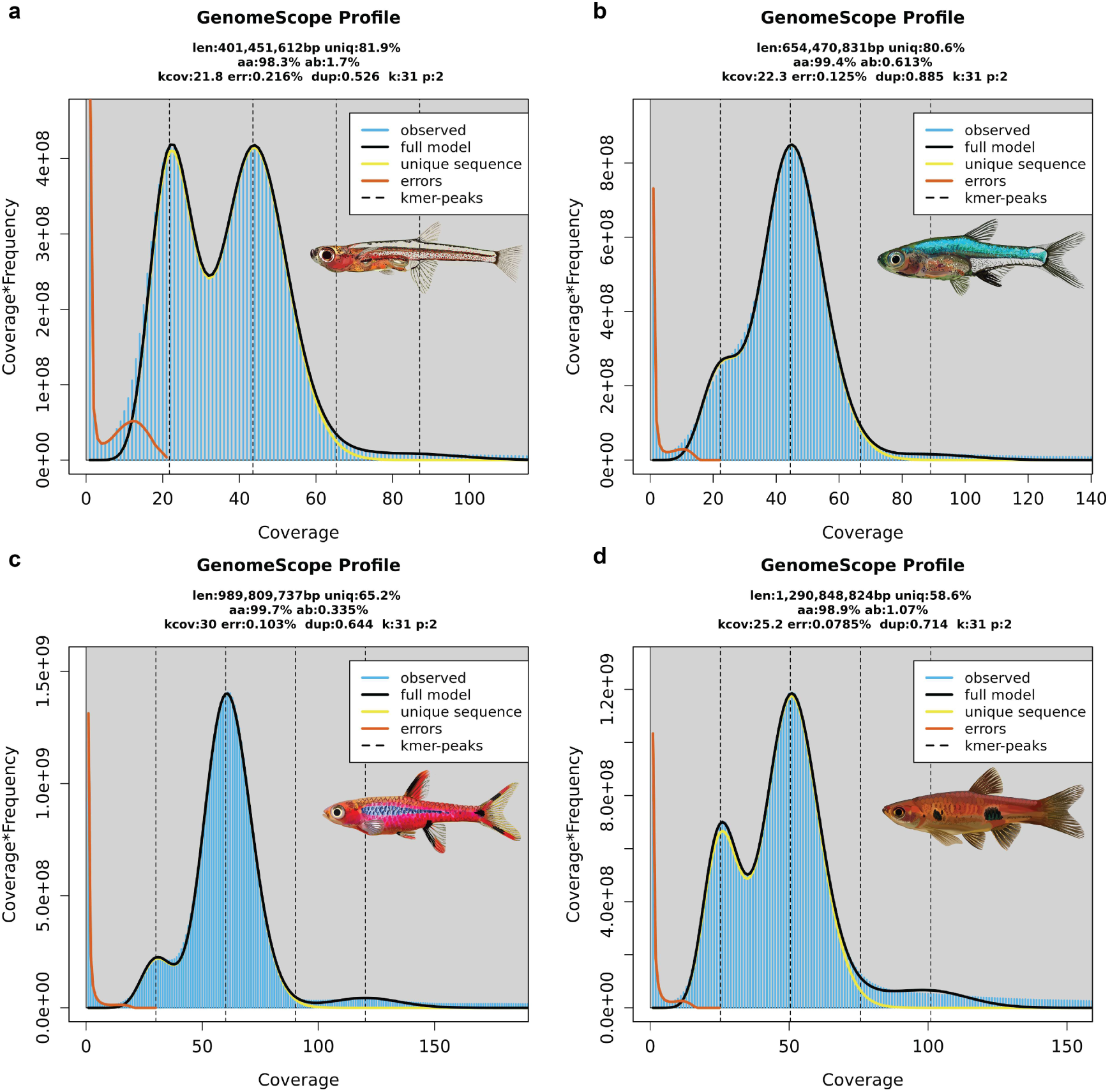
Frequency distributions of k-mers generated using GenomeScope2. Observed and modeled k-mer spectra are shown for each species, providing estimates of genome size, heterozygosity, and repeat content from unassembled PacBio HiFi reads: **(a)** *Paedocypris* sp.; **(b)** *Sundadanio atomus*; **(c)** *Boraras brigittae*; and **(d)** *Rasbora kalochroma*. The abbreviations at the top of each panel stands for genome size (len), unique k-mers (uniq), homozygosity (aa), heterozygosity (ab), haploid k-mer coverage (kcov), error rate (err), duplication rate (dup), k-mer used (k), and ploidy used (p).

### Genome assembly

The final assembly sizes were slightly larger than the k-mer–based genome size estimates for all four species (Supplementary Table 3). This could be expected as k-mer approaches typically underestimate repetitive content and the genome assemblies may retain alternative haplotypes in heterozygous regions, inflating the total assembly length (Pflug et al., 2020; Roach et al., 2018). The initial assemblies of *R. kalochroma* and *B. brigittae* were highly contiguous, consisting of 193 and 278 contigs with contig N50 values of 22.41 and 14.87 Mb, respectively. In contrast, *S. atomus* and *Paedocypris sp*. produced more fragmented initial assemblies, containing 913 and 1768 contigs and correspondingly smaller contig N50 values of 1.86 and 1.61 Mb (Supplementary Table 3). Integration of Hi-C data markedly improved contiguity, yielding scaffold-level assemblies of 91, 167, and 159 scaffolds with scaffold N50 values of 54.05, 41.49, and 30.49 Mb for *R. kalochroma*, *B. brigittae*, and *S. atomus*, respectively (Supplementary Table 3). For *Paedocypris* sp., Hi-C scaffolding resulted in 952 scaffolds with a scaffold N50 of 30.71 Mb (Supplementary Table 3). The snail plots in Fig. 3 provides a summary of the assembly statistics. The distribution of assembly scaffolds on coverage and GC proportion is shown in Supplementary Fig.1, while Supplementary Fig.2 illustrates the cumulative assembly plots. After manual curation, >98% of the genomes of *R. kalochroma*, *B. brigittae*, and *S. atomus* were anchored into pseudo-chromosomes (n = 25 for *R. kalochroma* and *B. brigittae*; n = 24 for *S. atomus*), whereas 88.1% of the *Paedocypris* sp. assembly was anchored into 15 pseudo-chromosomes (Supplementary Fig. 3), with the remaining sequence contained in small unplaced scaffolds (Supplementary Fig. 3a). The inferred chromosome numbers for *R. kalochroma*, *B. brigittae*, and *Paedocypris* sp. are consistent with published karyotypes for their respective genera (Aiumsumang et al., 2021; Donsakul et al., 2009; S. Liu et al., 2012). Although karyotypic data are unavailable for *Sundadanio*, the pseudo-chromosome count for *S. atomus* falls within the typical diploid range (n = 22–26) reported for cypriniform fishes (Arai, 2011). The GC content across assemblies ranged from 36.79% to 39.73%, lowest in *R. kalochroma* and highest in *Paedocypris* sp. (Supplementary Table 3). Merqury analyses indicated high consensus accuracy, with >99% k-mer completeness and QV scores above 50 for all four genome assemblies (Supplementary Fig. 4; Supplementary Table 3). BUSCO completeness using the actinopterygii_odb10 dataset exceeded 97.9% in *R. kalochroma*, *B. brigittae*, and *S. atomus*, and was 94.45% in *Paedocypris* sp., with duplicated BUSCOs accounting for <2.3% of genes in every assembly (Fig. 4; Supplementary Table 3). Collectively, these metrics demonstrate that all four assemblies exhibit high contiguity, accuracy, and completeness (Supplementary Table 3).

**Fig. 3.**
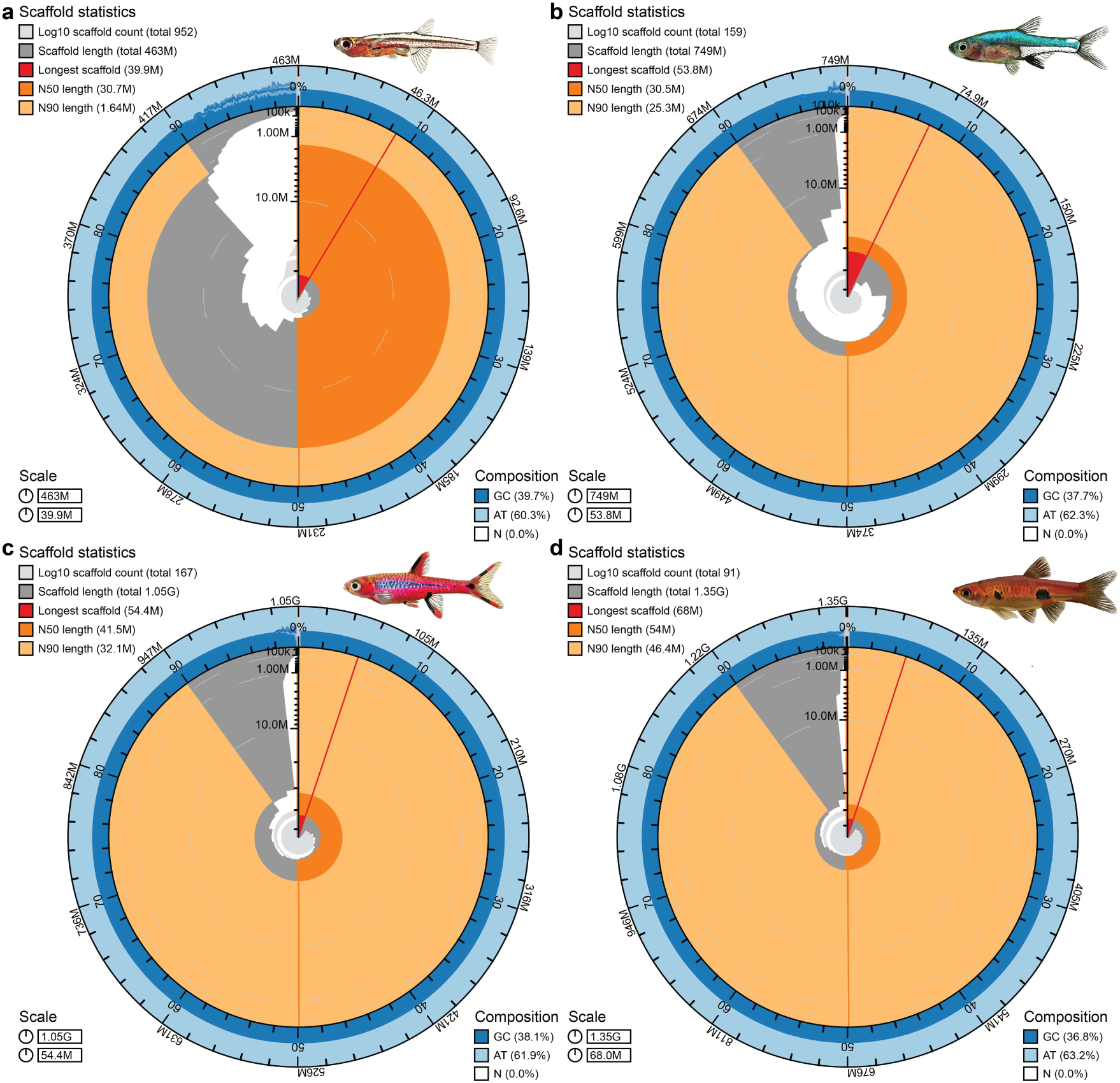
The BlobToolKit snail plots for the genome assemblies of **(a)** *Paedocypris* sp.; **(b)** *Sundadanio atomus*; **(c)** *Boraras brigittae*; and **(d)** *Rasbora kalochroma*. The circular plot is partitioned into 1,000 bins ordered by sequence size, each corresponding to 0.1% of the total assembly. Sequence length frequencies are shown in dark grey, with the radial scale normalized to the longest sequence in the assembly (highlighted in red). The N50 and N90 sequence lengths are indicated by orange and light-orange arcs, respectively. A light grey spiral represents the cumulative number of sequences plotted on a logarithmic scale, with white reference lines marking successive orders of magnitude. The outer blue and light-blue bands depict the distributions of GC, AT, and N content across the same bins used in the central plot.

**Fig. 4.**
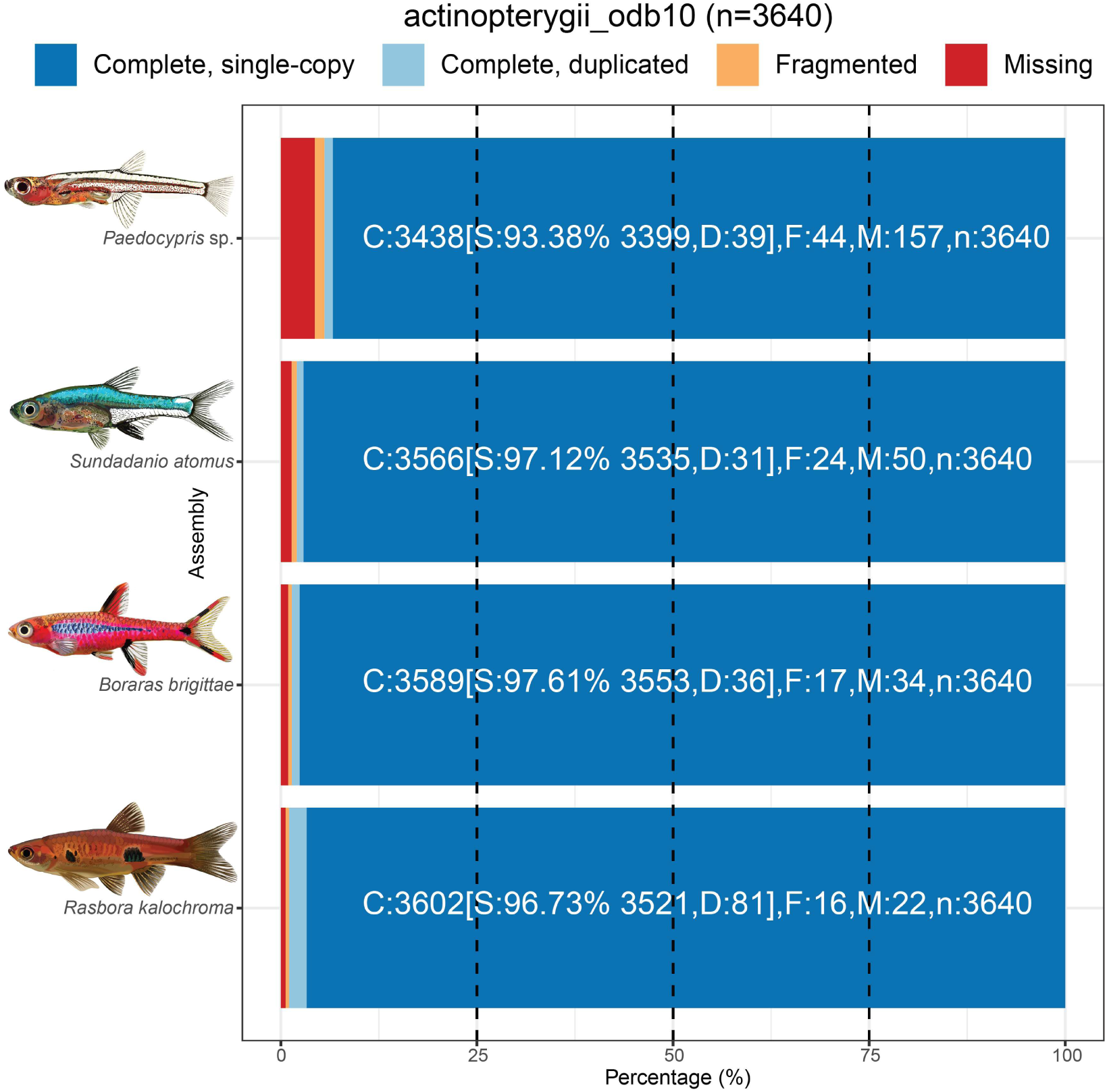
Summary of BUSCO genome completeness for *Paedocypris* sp., *Sundadanio atomus*, *Boraras brigittae*, and *Rasbora kalochroma* using the actinopterygii_odb10 dataset. Bars show complete, duplicated, fragmented, and missing orthologs.

Whole-genome dot-plot comparisons against the zebrafish GRCz11 reference (n=25 chromosomes) revealed extensive syntenic linearity for *R. kalochroma*, *B. brigittae*, and *S. atomus*, with the majority of their pseudo-chromosomes displaying one-to-one collinear relationships to the corresponding zebrafish chromosomes (Supplementary Fig. 5b-d). In contrast, the pseudo-chromosomes of *Paedocypris* sp. exhibited deviations from this pattern, showing reduced collinearity, and multiple disrupted syntenic blocks in comparison to the zebrafish reference (Supplementary Fig. 5a). This rearranged genomic architecture with the markedly reduced chromosome number in *Paedocypris*, could suggest that substantial lineage-specific rearrangements accompanied its extreme miniaturization and genome compaction.

### Annotation of repetitive elements

The proportion of each genome occupied by repetitive elements varied markedly among the four species (Fig. 5; Supplementary Table 4). The two progenetic miniatures, *Paedocypris* sp. and *S. atomus*, exhibited the lowest overall repeat content, spanning 35.2% and 41.5% of their genomes, respectively. In contrast, the proportioned dwarf *B. brigittae* and the non-miniature *R. kalochroma* displayed substantially higher repeat content (54.8% and 61.9%, respectively).

**Fig. 5.**
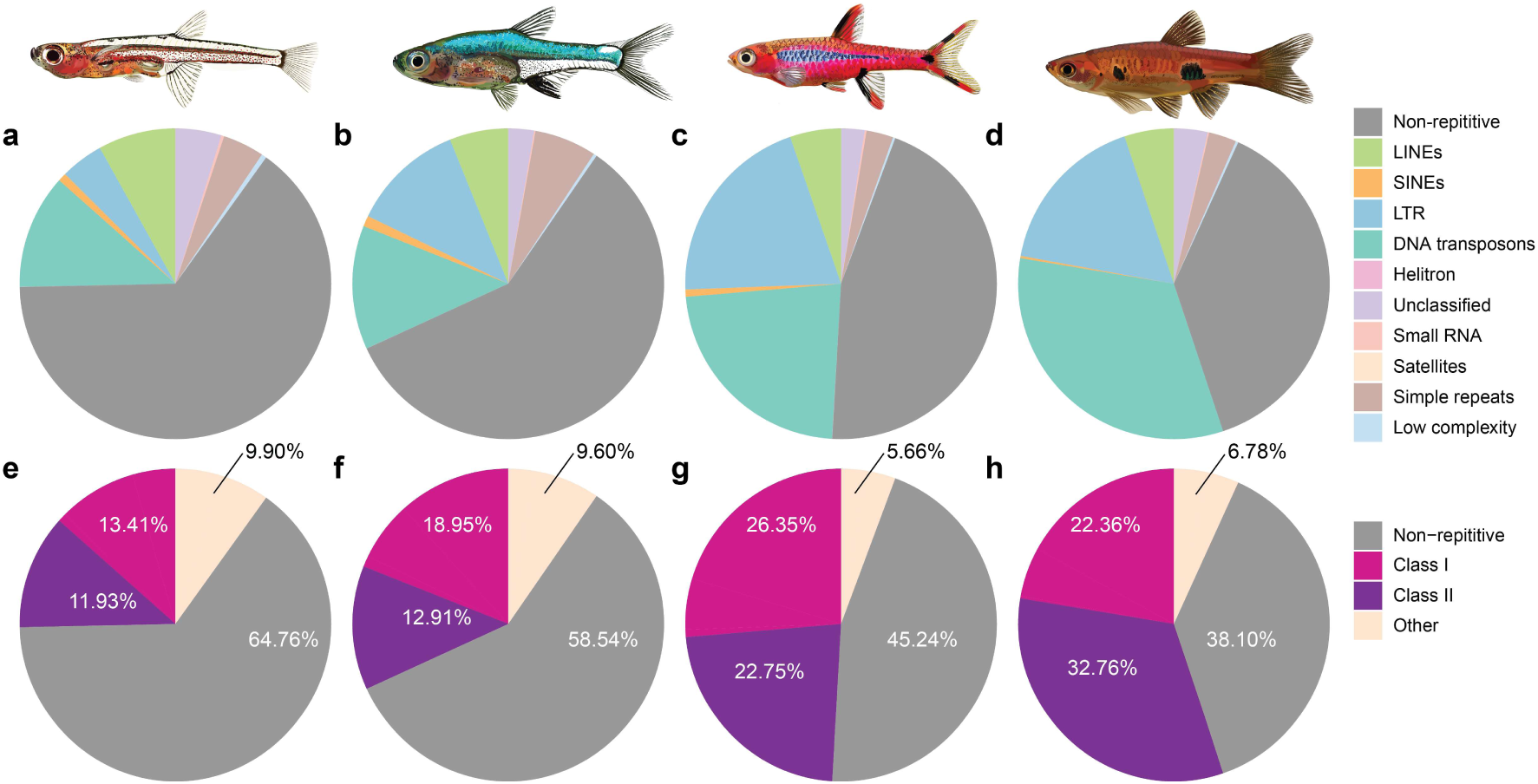
Proportion of the genome occupied by major repeat and transposable element categories in **(a,e)** *Paedocypris* sp., **(b,f)** *Sundadanio atomus*, **(c,g)** *Boraras brigittae*, and **(d,h)** *Rasbora kalochroma*.

Differences in the relative abundance of major repeat classes further distinguish the progenetic miniatures from the other two species (Fig. 5; Supplementary Table 4). Class I retrotransposons were present at broadly comparable levels across all species, ranging from 13.41% in *Paedocypris* sp. to 26.35% in *B. brigittae*. By contrast, Class II DNA transposons showed lineage-specific variation: *B. brigittae* and *R. kalochroma* exhibited substantial expansions (22.75% and 32.76%, respectively), whereas *Paedocypris* sp. and *S. atomus* contained much lower proportions (11.93% and 12.91%, respectively). These differences were largely driven by DNA transposons themselves, which dominate the Class II fraction in all four species (Fig. 5; Supplementary Table 4).

In contrast to this pattern, the category comprising “other repeats” including small RNAs, satellites, simple repeats, low-complexity regions, and unclassified elements was more abundant in the two progenetic miniatures (9.90% in *Paedocypris* sp. and 9.60% in *S. atomus*) than in *B. brigittae* (5.66%) and *R. kalochroma* (6.78%). This trend was mainly influenced by simple repeats, which were elevated in *Paedocypris* sp. and *S. atomus* (4.3% and 6.45%) relative to the other two species (2.79% and 2.87%: Fig. 5; Supplementary Table 4). Additionally, *Paedocypris* sp. possessed the lowest proportion of LTR elements (4.45%) and the highest proportion of unclassified repeats (4.78%) among the four species (Fig. 5; Supplementary Table 4).

Taken together, these patterns show that while total genome size broadly correlates with overall repeat content, the evolutionary trajectories of TE expansion is different among lineages. Interestingly, the species with the two smallest genomes (*Paedocypris* sp. and *S. atomus*) do not show the smallest TE expansions across all the different TE types, whereas *B. brigittae* and *R. kalochroma*, with larger genomes, show disproportionately large expansions of Class II DNA transposons. This could suggest that overall genome size alone does not predict the scale or direction of TE proliferation (Feschotte & Pritham, 2007; Kapusta et al., 2017; Sotero-Caio et al., 2017).

Repeat landscapes revealed further differences in TE dynamics among the four species (Fig. 6, Supplementary Fig. 6). Repeat landscapes plot the distribution of TE copies as a function of Kimura substitution levels, which reflect divergence from consensus TE sequences and therefore approximate the temporal profile of transpositional activity (Rodriguez C Arkhipova, 2023). *Boraras brigittae* and *R. kalochroma* displayed strong peaks at low Kimura distances (Fig. 6, Supplementary Fig. 6), consistent with substantial recent TE activity and ongoing bursts of transposition (Sotero-Caio et al., 2017). In contrast, the progenetic miniatures exhibited bimodal landscapes characterized by an accumulation of older, highly diverged elements and comparatively little evidence of recent activity. In *Paedocypris* sp., the secondary peak was dominated by “other repeat” categories, rather than Class I or Class II repeat classes (Fig. 6a).

**Fig. 6.**
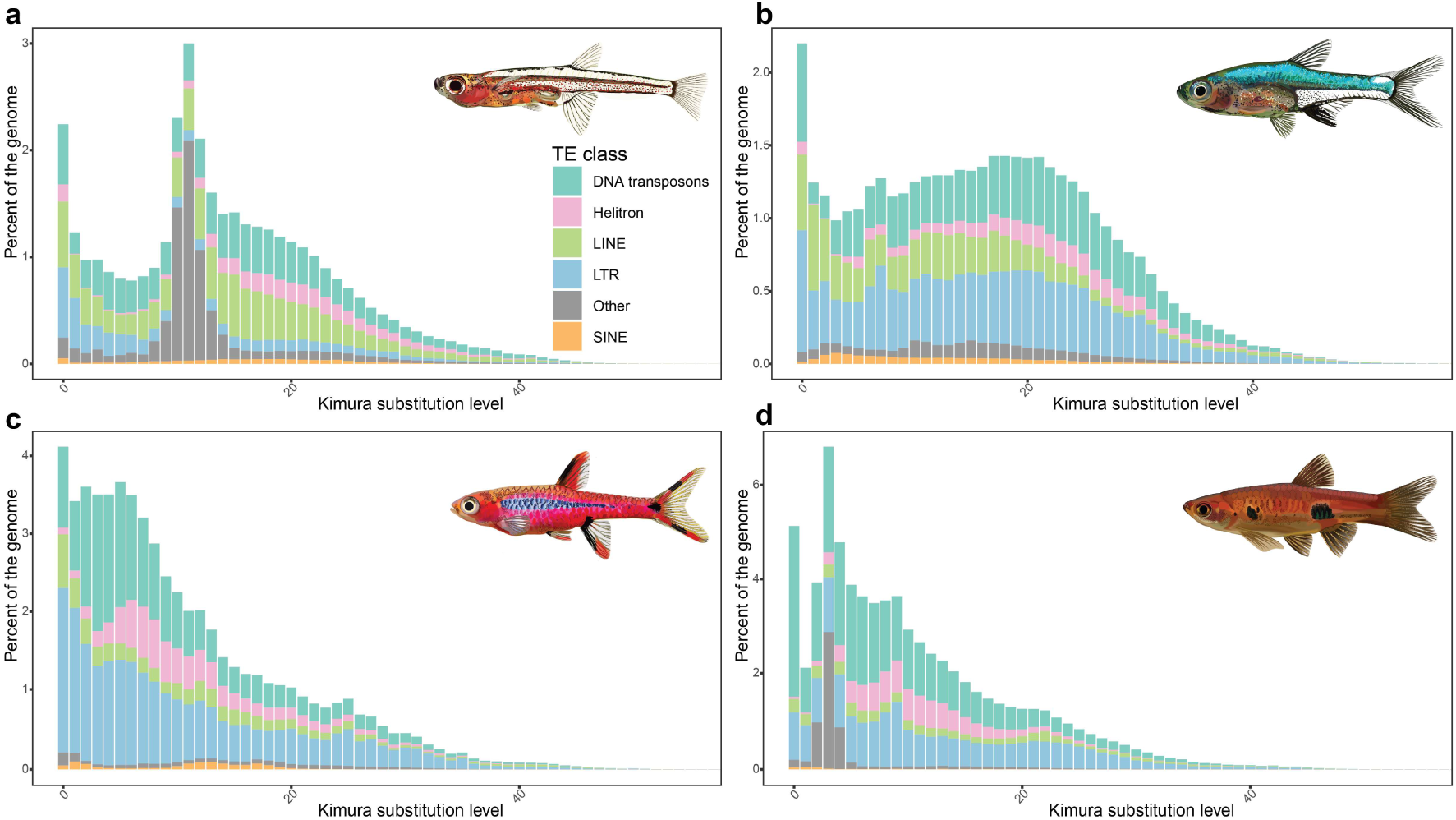
Repeat landscapes for major transposable element (TE) subclasses and additional repeat categories (“other” repeats: small RNAs, satellites, simple repeats, low-complexity regions, and unclassified elements) in **(a)** *Paedocypris* sp., **(b)** *Sundadanio atomus*, **(c)** *Boraras brigittae*, and **(d)** *Rasbora kalochroma*. The x-axis displays CpG-adjusted Kimura substitution levels (%), representing divergence from TE consensus sequences. Low-divergence peaks (left) indicate recent or ongoing TE activity, whereas high-divergence peaks (right) reflect ancient insertions.

### Annotation of protein coding genes

BRAKER3 yielded 28428, 20801, 22180, and 35129 mRNA predictions in *S. atomus*, *B. brigittae*, *Paedocypris sp*., and *R. kalochroma*, respectively, of which 23107, 17206, 18608, and 29670 were designated as gene models (Supplementary Table 5).

Mean gene length differed among species, with longer genes in *B. brigittae* (16692 bp) and *R. kalochroma* (19745 bp) relative to *S. atomus* (12079 bp) and *Paedocypris* sp. (9314 bp). Despite this, coding-region structure was highly conserved: mean CDS length (1552-1667 bp), mean exon length (168-175 bp), and exon count per mRNA (8.9-10.0) were similar across all four genomes (Supplementary Table 5).

The primary driver of gene-length differences was intron architecture. Mean intron lengths were substantially longer in *B. brigittae* (1966 bp) and *R. kalochroma* (2355 bp) compared with *S. atomus* (1286 bp) and *Paedocypris* sp. (875 bp). This pattern extends the findings of Malmstrøm et al. (2018), who reported pronounced intron reduction in *Paedocypris*. This could suggest that intron shortening may represent a more general feature of genome compaction in progenetic miniature cypriniform fishes. Pufferfishes, which exhibit some of the most compact genomes among vertebrates, also show pronounced intron shortening (K. Liu et al., 2025).

Gene-set completeness was high across species. BUSCO scores (actinopterygii_odb10) exceeded 95.2% for *R. kalochroma*, *B. brigittae*, and *S. atomus*, and reached 90.7% in *Paedocypris* sp. with duplicated BUSCOs accounting for <3.9% in all the gene annotations (Supplementary Table 5). OMArk assessments likewise indicated robust proteome quality, with 80.0-96.8% completeness across 16205 hierarchical orthologous groups and >92.8% internal consistency, with no contamination and minimal fragmentary mappings (Fig. 7; Supplementary Table 5). These values are comparable to other high-quality cypriniform proteomes curated in the OMArk database.

**Fig. 7.**
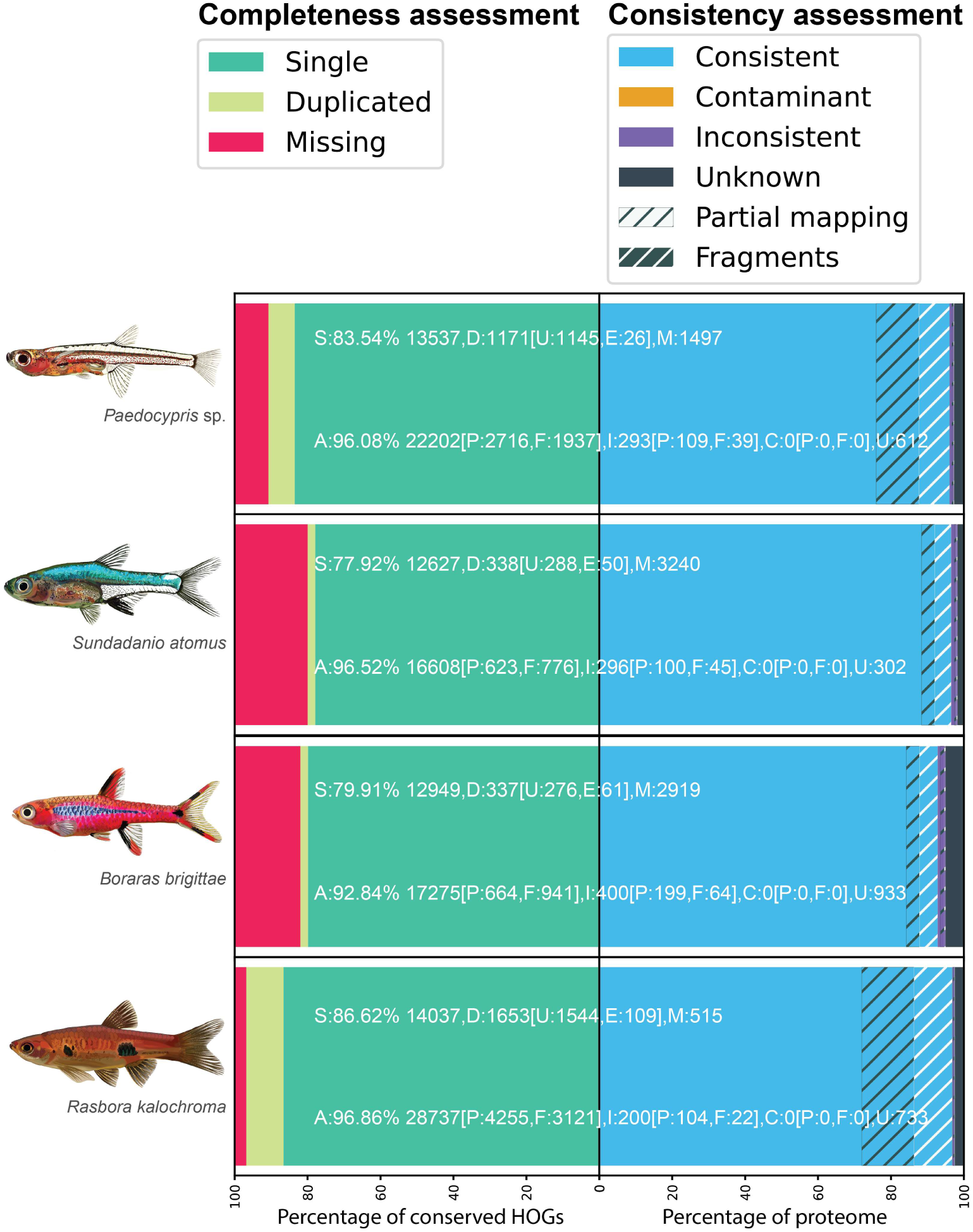
OMArk completeness and consistency metrics for the predicted proteomes of *Paedocypris* sp., *Sundadanio atomus*, *Boraras brigittae*, and *Rasbora kalochroma*. Completeness reflects recovery of expected ancestral gene set of the species’ lineage; consistency indicates whether proteins map to the correct lineage, an incorrect lineage, show contamination, or remain unassigned.

## Conclusions

In this study, we present highly contiguous and complete genomes representing two progenetic miniatures (*Paedocypris* sp. and *S. atomus*), a proportioned dwarf (*B. brigittae*), and a non-miniature species (*R. kalochroma*) which are the first reference genome assemblies for freshwater fishes inhabiting the acidic blackwaters of Southeast Asia’s peat-swamp forests, an extreme and understudied ecosystem. Importantly, the assemblies include a reference genome for *Paedocypris*, one of the smallest vertebrates known, offering new opportunities to investigate the genomic basis of extreme miniaturization.

Together, these genomes establish a comparative framework for dissecting how independent cypriniform lineages have adapted to the physiologically challenging conditions of highly acidic, oxygen-depleted, and ion-poor blackwaters. They also provide a foundation for exploring broader questions in vertebrate evolution, including the genomic correlates of progenesis, developmental truncation, and repeated miniaturization. Furthermore, the phylogenetic placement of *Paedocypris* and *Sundadanio* within Cypriniformes has remained recalcitrant in sequence-based phylogenomic analyses (Sudasinghe et al., 2026). The availability of chromosome-level genome assemblies for both genera now enables the evaluation of their evolutionary relationships using complementary genomic characters, including rare genomic changes and conserved patterns of gene synteny (Steenwyk & King, 2024). Beyond evolutionary and developmental biology, these genomic resources hold value for conservation, enabling assessments of evolutionary distinctiveness and adaptive potential in one of the most threatened freshwater habitats in the world.

## Acknowledgments

We thank Pamela Nicholson and the staff of the Next Generation Sequencing (NGS) Platform at the University of Bern, Switzerland, for their dedicated support in data generation, as well as for their constructive discussions and invaluable troubleshooting, and Nadeela Hirimuthugoda (https://nadeela.weebly.com/illustrations.html) for providing the fish illustrations.

## Funding

The study was funded by the Swiss National Science Foundation (grant 310030_185120).

---

**Supplementary Fig. 1.**
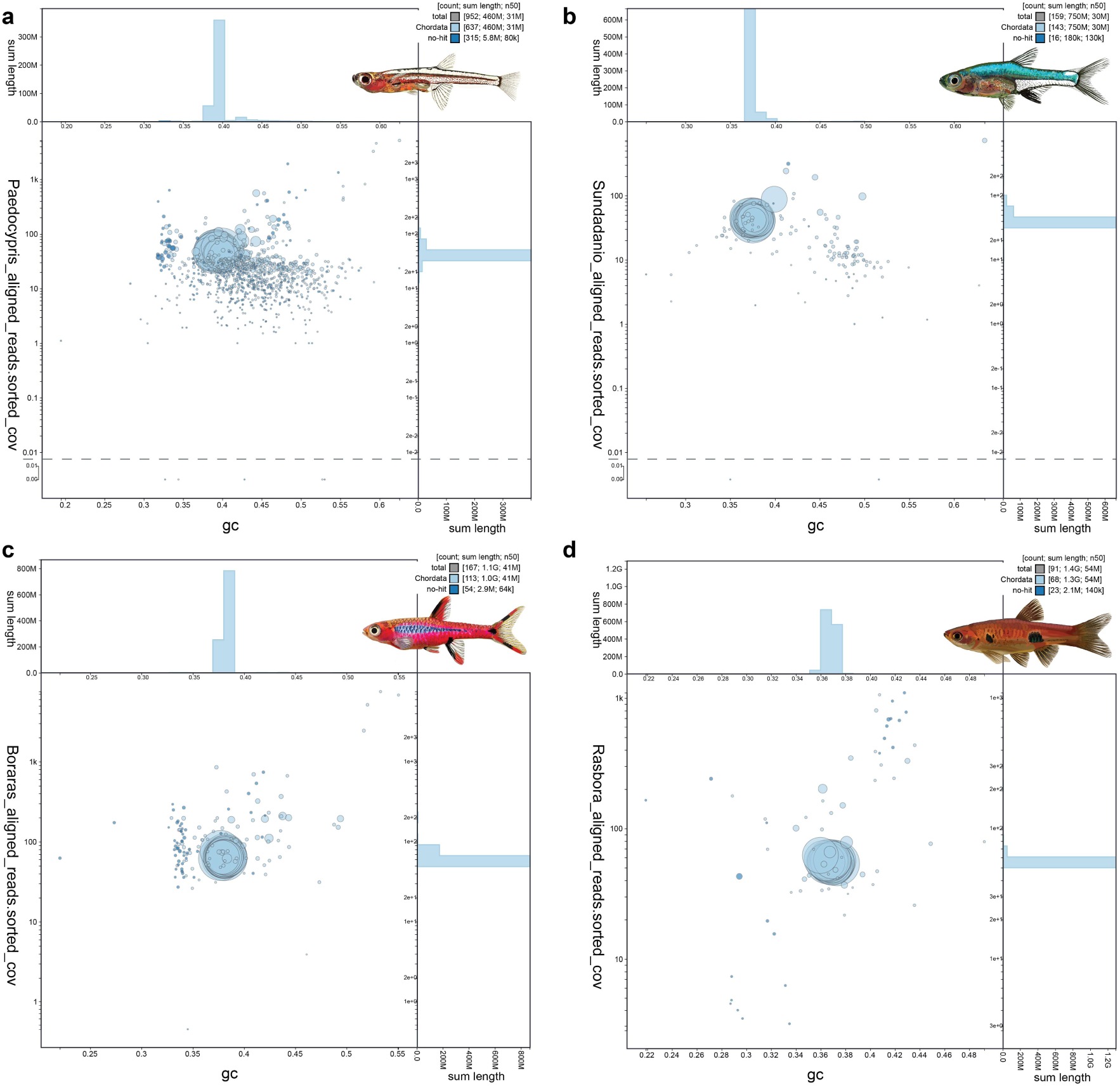
BlobToolKit GC-coverage plot of **(a)** *Paedocypris* sp.; **(b)** *Sundadanio atomus*; **(c)** *Boraras brigittae*; and **(d)** *Rasbora kalochroma*. Scaffolds are colour-coded according to phylum, with circle sizes scaled to scaffold length. Histograms along each axis display the distribution of cumulative scaffold lengths.

**Supplementary Fig. 2.**
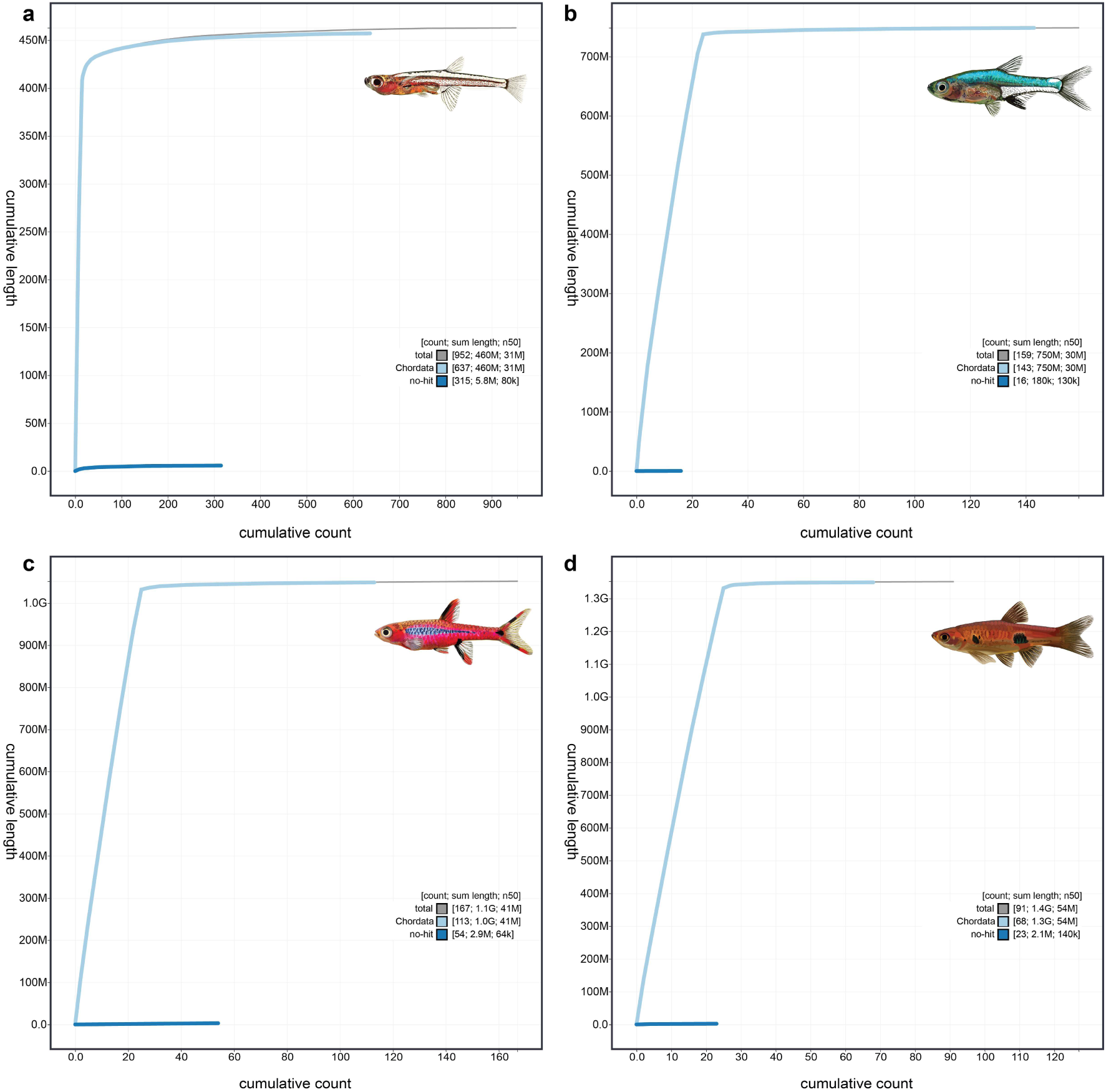
BlobToolKit cumulative sequence plot of **(a)** *Paedocypris* sp.; **(b)** *Sundadanio atomus*; **(c)** *Boraras brigittae*; and **(d)** *Rasbora kalochroma*. The grey curve represents the cumulative length of all scaffolds, while coloured curves indicate the cumulative lengths of scaffolds assigned to individual phyla.

**Supplementary Fig. 3.**
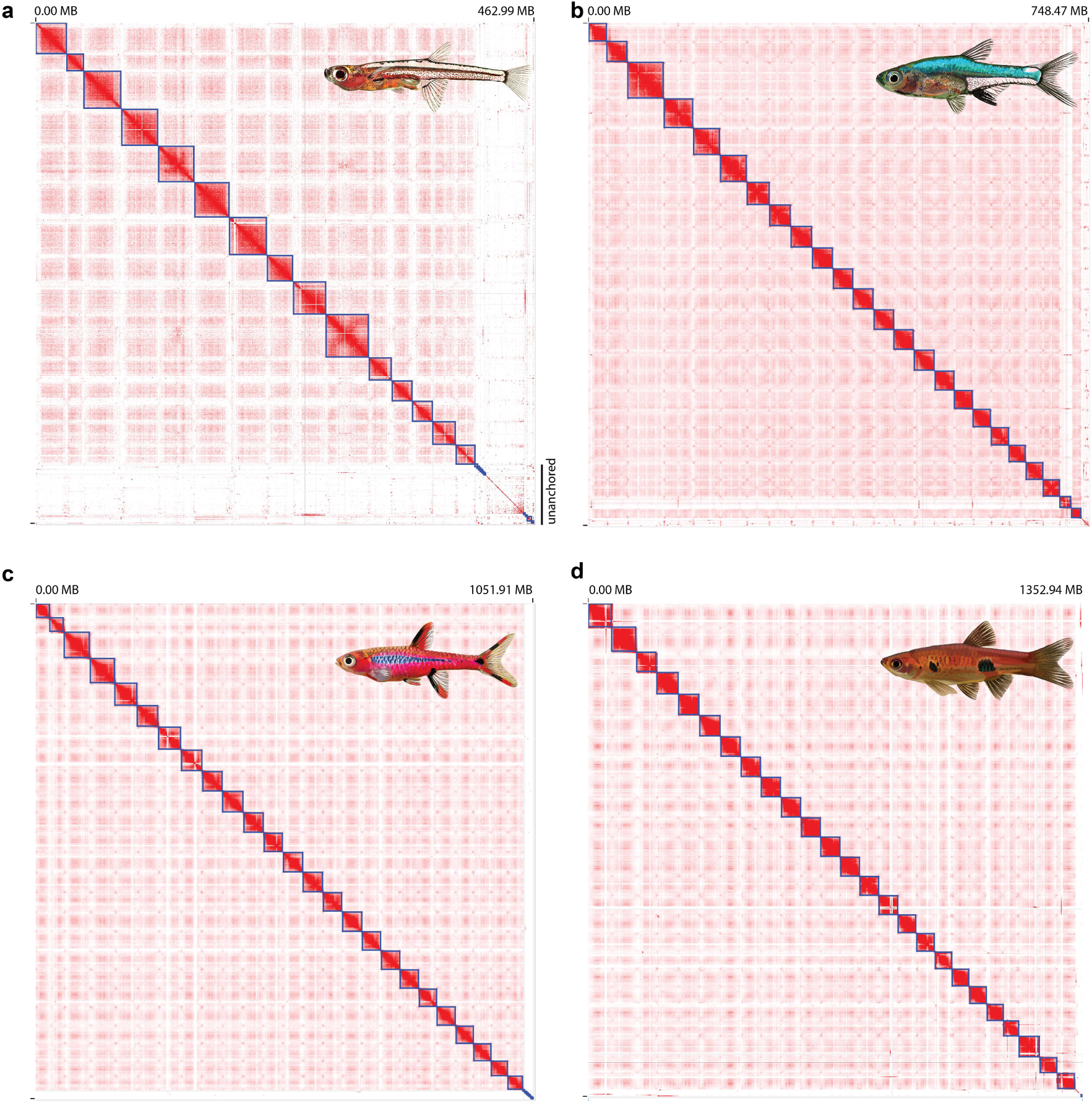
Hi-C interaction heatmaps generated with Juicebox for **(a)** *Paedocypris* sp.; **(b)** *Sundadanio atomus*; **(c)** *Boraras brigittae*; and **(d)** *Rasbora kalochroma*. Darker blocks indicate stronger interaction signals.

**Supplementary Fig. 4.**
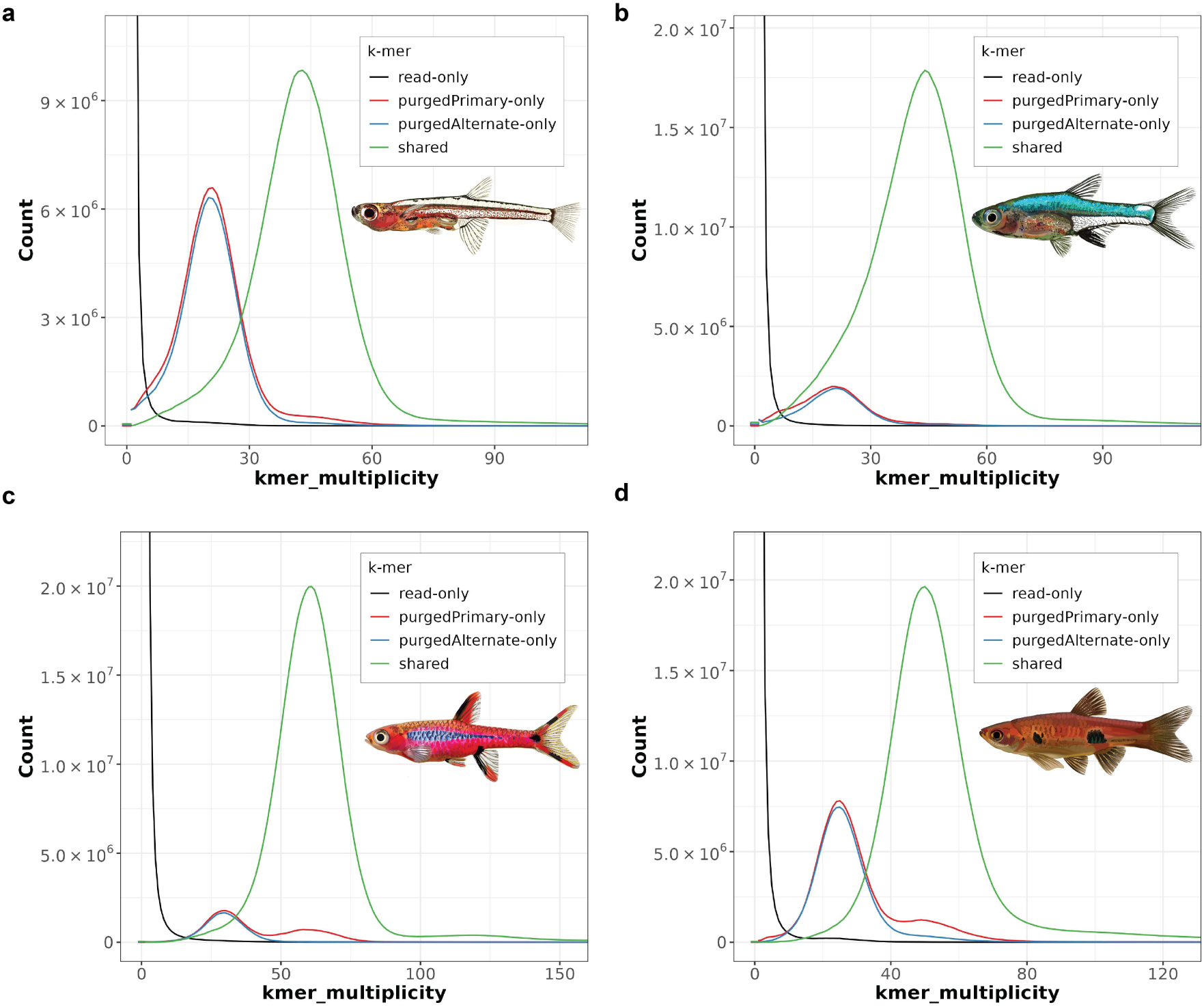
Merqury k-mer spectra plots for **(a)** *Paedocypris* sp., **(b)** *Sundadanio atomus*, **(c)** *Boraras brigittae*, and **(d)** *Rasbora kalochroma*. These plots depict the recovery of sequencing read k-mers in the final assemblies, providing an assessment of completeness and haplotype representation. The horizontal axis indicates k-mer multiplicity and the vertical axis the number of k-mers. Black curves represent k-mers present in the read set but absent from the assembly. Green peaks correspond to homozygous k-mers shared between haplotypes, whereas red and blue peaks indicate heterozygous k-mers unique to each haplotype.

**Supplementary Fig. 5.**
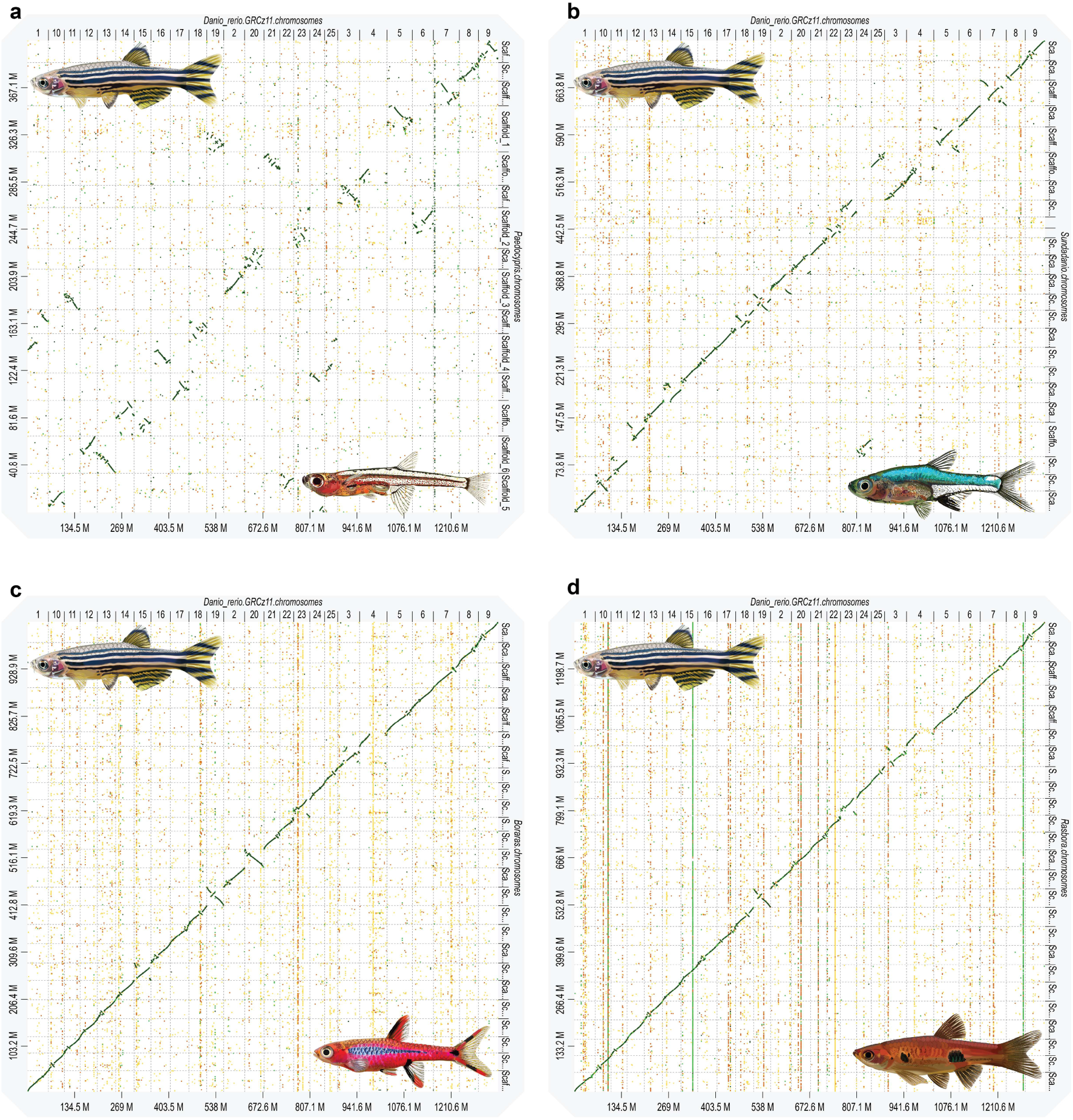
Dot plot comparisons of **(a)** *Paedocypris* sp., **(b)** *Sundadanio atomus*, **(c)** *Boraras brigittae*, and **(d)** *Rasbora kalochroma* genome assemblies against the *Danio rerio* GRCz11 reference genome, generated using D-GENIES. These plots illustrate large-scale synteny and chromosomal rearrangements relative to zebrafish.

**Supplementary Fig. 6.**
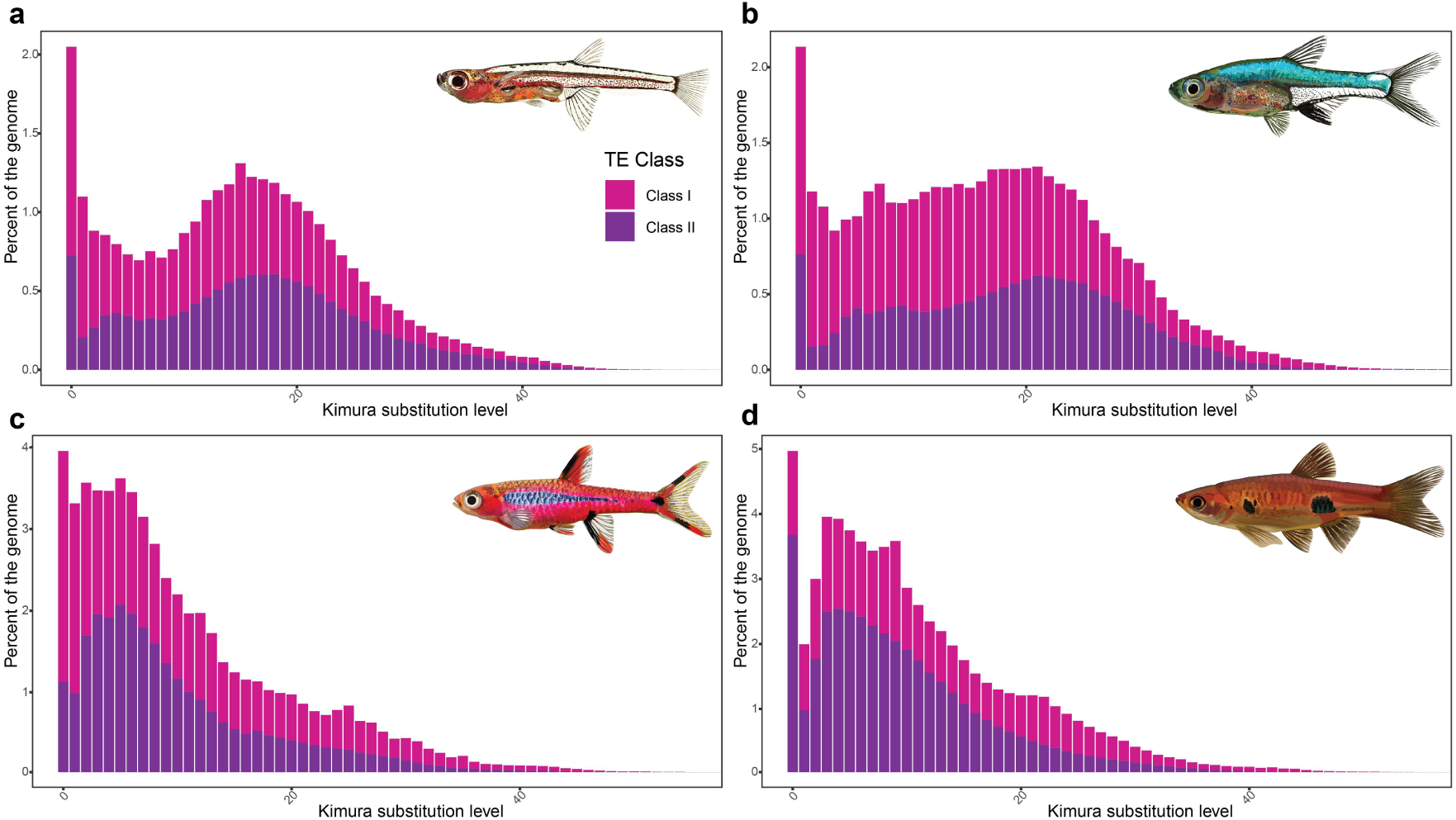
Repeat landscapes for Class I and Class II transposable elements in **(a)** *Paedocypris* sp., **(b)** *Sundadanio atomus*, **(c)** *Boraras brigittae*, and **(d)** *Rasbora kalochroma*. The x-axis shows CpG-adjusted Kimura substitution levels (%), representing divergence from TE consensus sequences. Elements with low divergence (left side) correspond to recently active TE insertions, whereas those with higher divergence (right side) reflect older transposition events.

**Supplementary Table 1.** Sample and sequencing information.

| Metadata | *Paedocypris* sp. | *Sundadanio atomus* | *Boraras brigittae* | *Rasbora kalochroma* |
| --- | --- | --- | --- | --- |
| BioProject |  |  |  |  |
| Specimen ID: PacBio HiFi | LR15863 | LR15870 | LR15871 | LR15877 |
| BioSample (source individual): PacBio HiFi |  |  |  |  |
| BioSample (tissue): PacBio HiFi |  |  |  |  |
| PacBio HiFi platform | PacBio Sequel IIe | PacBio Revio | PacBio Revio | PacBio Revio |
| Run accessions: PacBio HiFi |  |  |  |  |
| Read count total: PacBio HiFi | 3.57 million | 2.13 million | 4.68 million | 6.03 million |
| Base count total: PacBio HiFi | 18.79 Gb | 30.34 Gb | 61.55 Gb | 66.94 Gb |
| Specimen ID: Hi-C | LR15864 | LR15868 | LR15873 | LR15880 |
| BioSample (source individual): Hi-C |  |  |  |  |
| BioSample (tissue): Hi-C |  |  |  |  |
| Hi-C platform | Illumina NovaSeq 6000 | Illumina NovaSeq 6000 | Illumina NovaSeq 6000 | Illumina NovaSeq 6000 |
| Run accessions: Hi-C |  |  |  |  |
| Read count total: Hi-C | 111.47 million pairs | 202.84 million pairs | 204.23 million pairs | 330.66 million pairs |
| Base count total: Hi-C | 33.66 Gb | 61.25 Gb | 61.67 Gb | 99.86 Gb |
| Specimen ID: Iso-Seq | LR15860, LR15865 | LR15869 | LR15875 | LR15878 |
| BioSample (source individual): Iso-Seq |  |  |  |  |
| BioSample (tissue): Iso-Seq |  |  |  |  |
| Iso-Seq platform | PacBio Sequel IIe | PacBio Sequel IIe | PacBio Sequel IIe | PacBio Sequel IIe |
| Run accessions: Iso-Seq |  |  |  |  |
| Read count total: Iso-Seq | 1.73, 1.87 million | 1.48 million | 1.37 million | 1.42 million |
| Base count total: Iso-Seq | 1.49, 1.59 Gb | 2.81 Gb | 2.29 Gb | 2.47 Gb |
| Remarks | Progenetic miniature | Progenetic miniature | Proportioned dwarf | Non-miniature |

**Supplementary Table 2.**
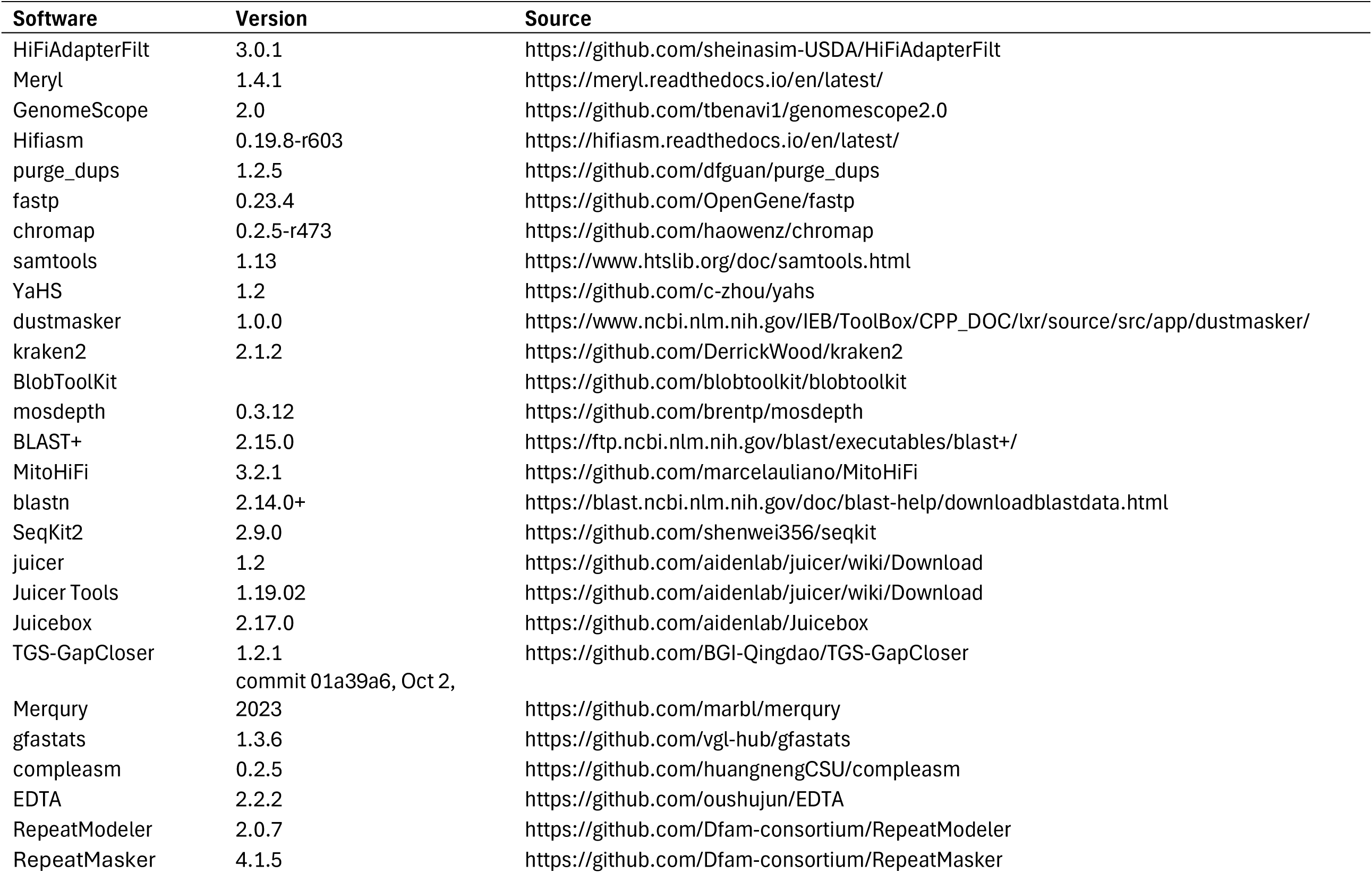
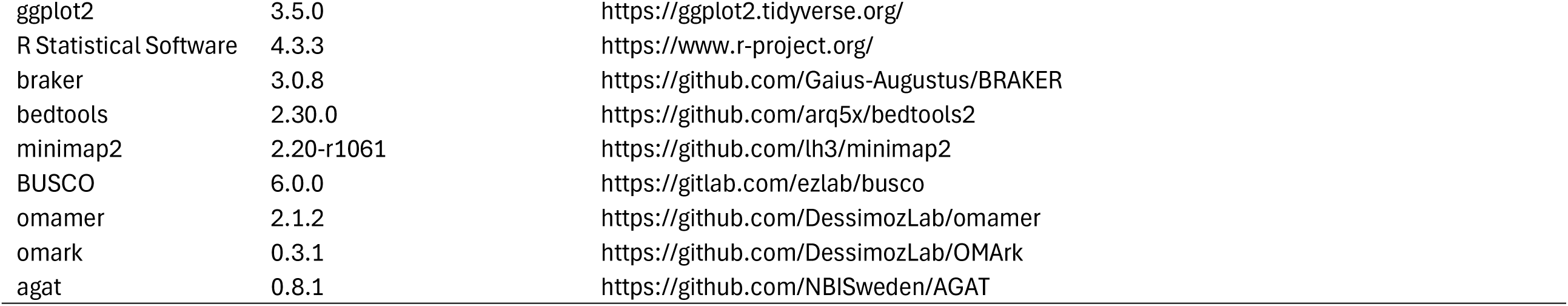
Software versions and sources employed in this study.

**Supplementary Table 3.**
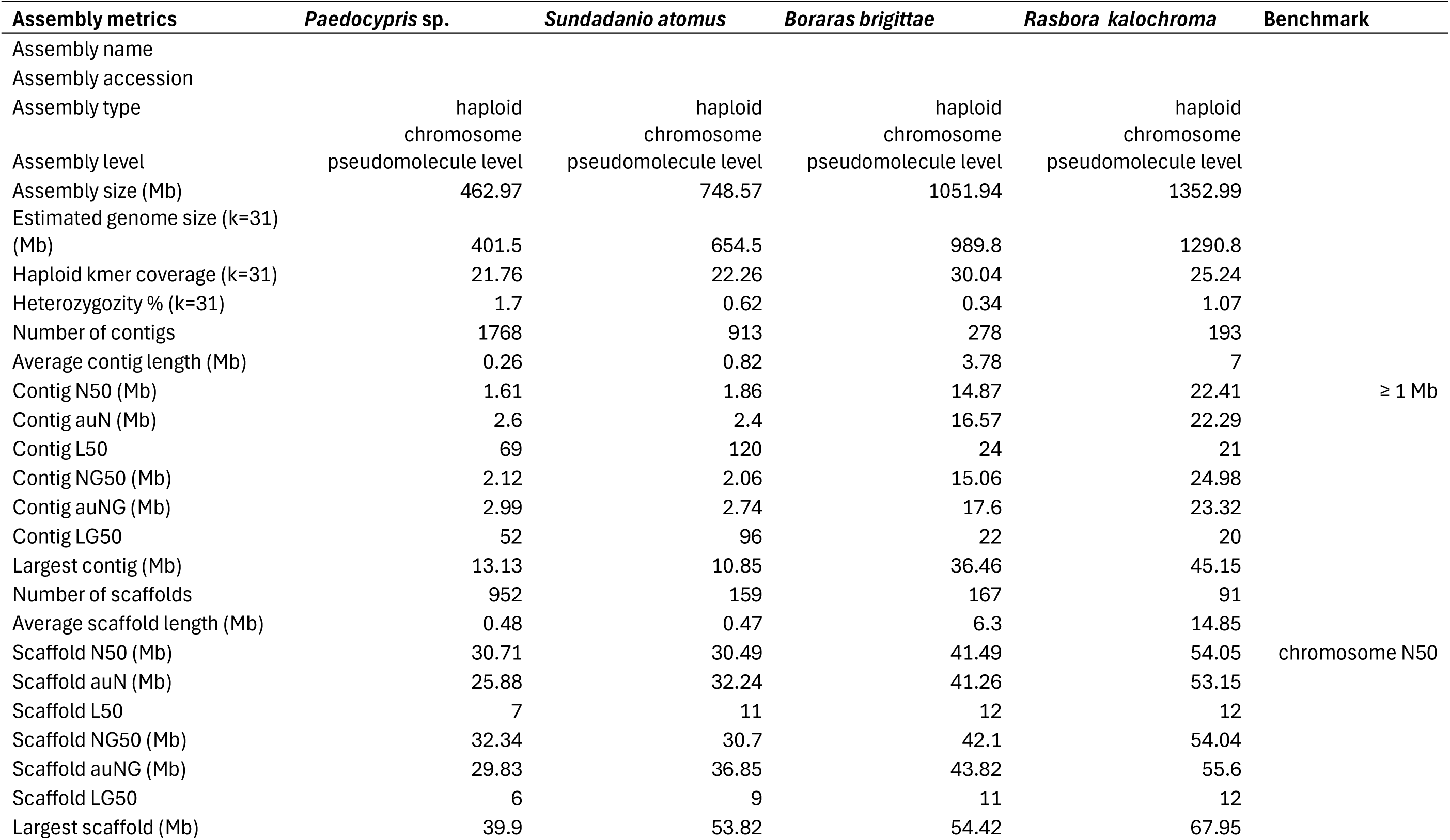
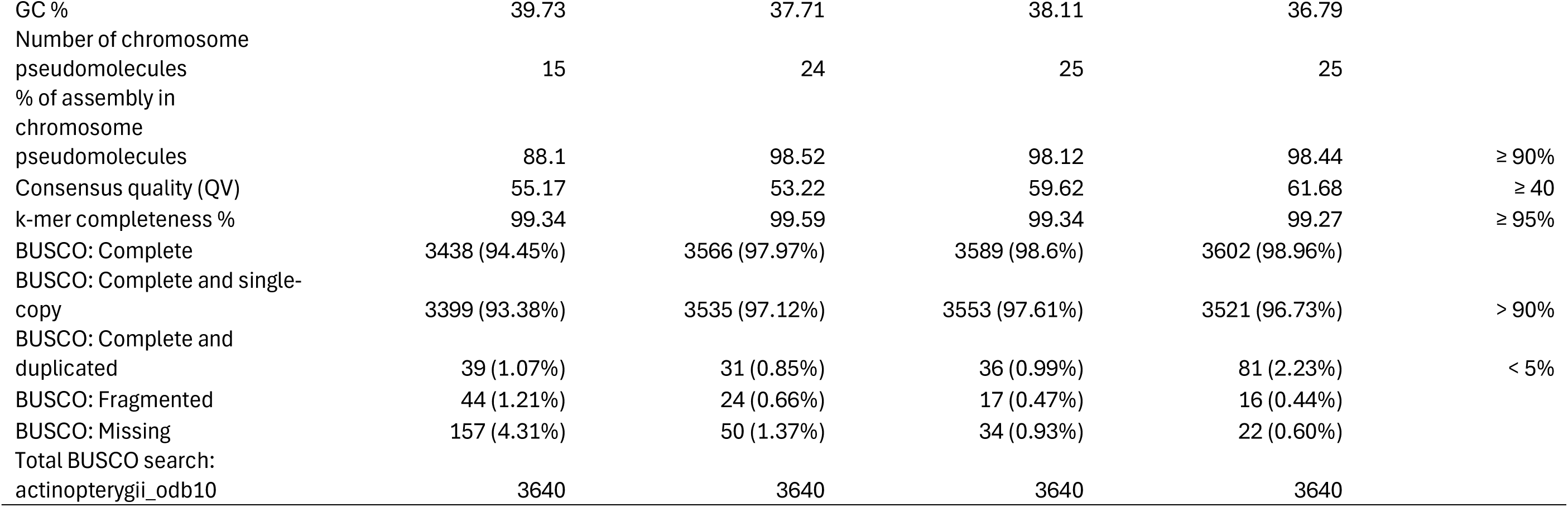
Genome assembly statistics.

**Supplementary Table 4.**
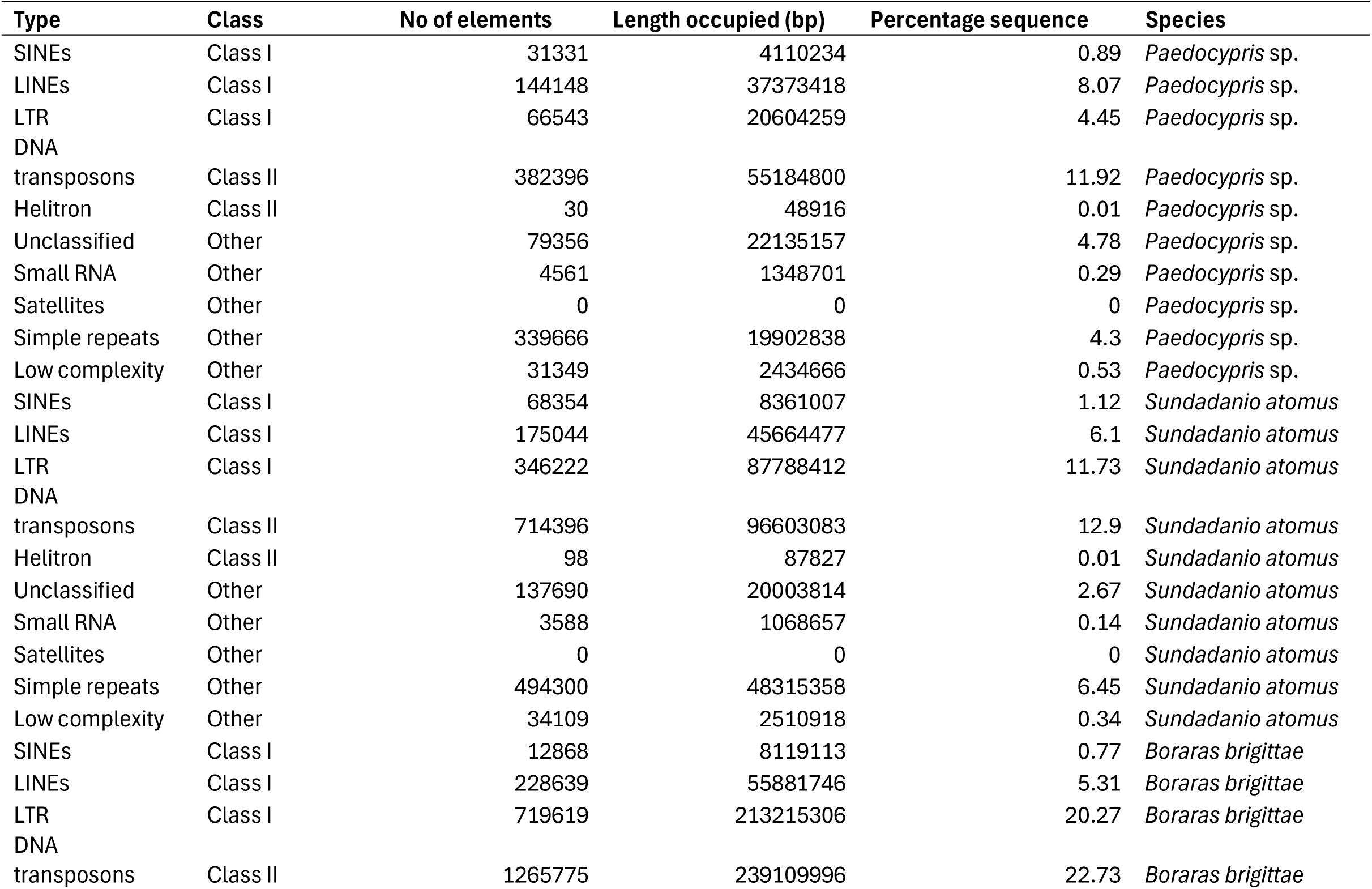
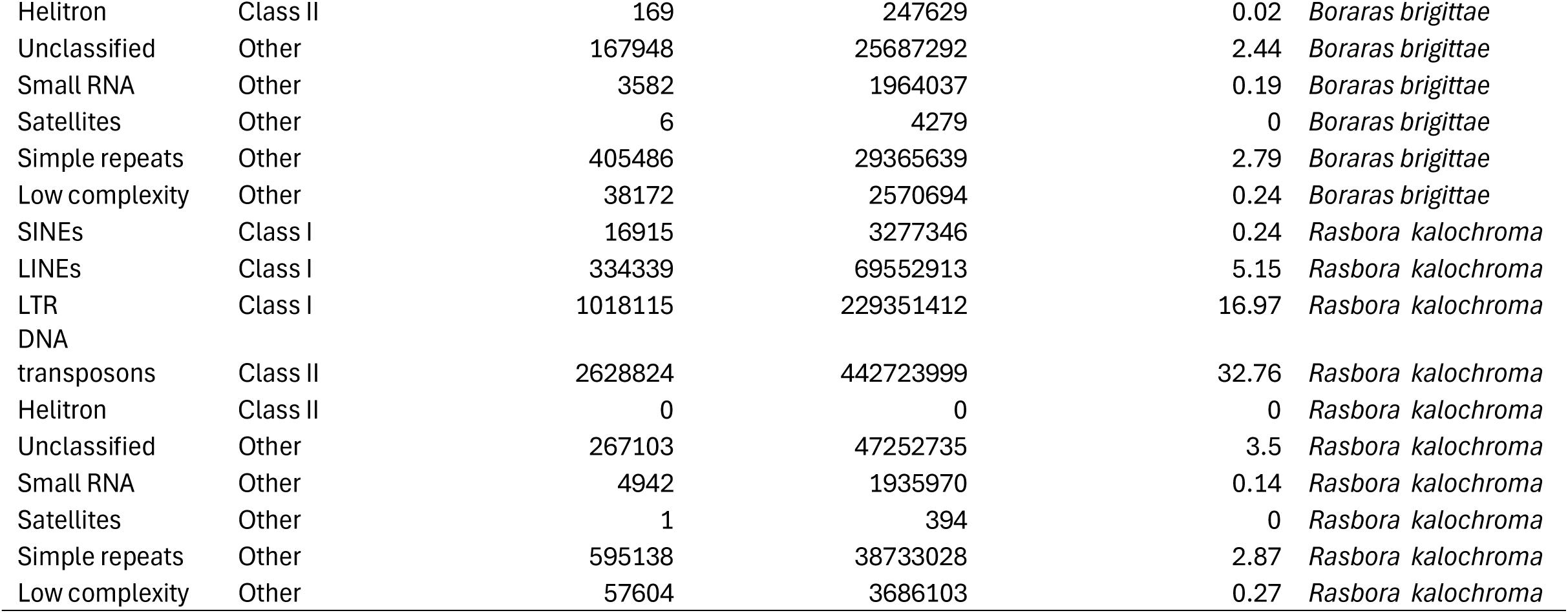
Proportion of each genome occupied by major repeat and transposable element (TE) categories. Note: RepeatMasker counts most repeats fragmented by insertions or deletions as a single element.

**Supplementary Table 5.** Gene annotation statistics.

| Annotation metrics | *Paedocypris* sp. | *Sundadanio atomus* | *Boraras brigittae* | *Rasbora kalochroma* |
| --- | --- | --- | --- | --- |
| Number of genes | 23107 | 17206 | 18608 | 29670 |
| Number of mRNAs | 28428 | 20801 | 22180 | 35129 |
| Mean gene length (bp) | 9314 | 12079 | 16692 | 19745 |
| Mean CDS length (bp) | 1667 | 1588 | 1552 | 1556 |
| Mean exon length (bp) | 169 | 168 | 171 | 175 |
| Mean intron length (bp) | 875 | 1286 | 1966 | 2355 |
| Mean exons per mRNA | 10 | 9.4 | 9.1 | 8.9 |
| BUSCO: Complete | 3301 (90.7%) | 3485 (95.7%) | 3515 (96.6%) | 3465 (95.2%) |
| BUSCO: Complete and single-copy | 3182 (87.4%) | 3396 (93.3%) | 3426 (94.1%) | 3322 (91.3%) |
| BUSCO: Complete and duplicated | 119 (3.3%) | 89 (2.4%) | 89 (2.4%) | 143 (3.9%) |
| BUSCO: Fragmented | 87 (2.4%) | 57 (1.6%) | 45 (1.2%) | 78 (2.1%) |
| BUSCO: Missing | 252 (6.9%) | 98 (2.7%) | 80 (2.2%) | 97 (2.7%) |
| Total BUSCO search | 3640 | 3640 | 3640 | 3640 |
| OMArk: Complete | 14708(90.77%) | 12965 (80.01%) | 13286 (81.99%) | 15690 (96.82%) |
| OMArk: Complete and single-copy | 13537 (83.54%) | 12627 (77.92%) | 12949 (79.91%) | 14037 (86.62%) |
| OMArk: Complete and duplicated | 1171 (7.23%) | 338 (2.09%) | 337 (2.08%) | 1653 (10.20%) |
| OMArk: Missing | 1497 (9.24%) | 3240 (19.99%) | 2919 (18.01%) | 515 (3.18%) |
| Total OMArk conserved HOGs | 16205 | 16205 | 16205 | 16205 |
| OMArk: Consistent | 22202 (96.08%) | 16608 (96.52%) | 17275 (92.84%) | 28737 (96.86%) |
| OMArk: Inconsistent | 293 (1.27%) | 296 (1.72%) | 400 (2.15%) | 200 (0.67%) |
| OMArk: Contaminant | 0 | 0 | 0 | 0 |
| OMArk: Unknown | 612 (2.65%) | 302 (1.76%) | 933 (5.01%) | 733 (2.47%) |
| OMArk: Proteome percentage total | 97.35% | 98.24% | 94.99% | 97.53% |

## Notes

### Competing Interest Statement

The authors have declared no competing interest.

